# An adaptive noradrenergic–prefrontal circuit for innate avoidance of heights

**DOI:** 10.64898/2026.09.14.751402

**Authors:** Stephanie M. Staszko, Emi Krishnamurthy, Rohan Lokanadham, Jessica P. He, Aakash Basu, Jocelyne Rondeau, Abigail L. Yu, John H. Krystal, Alfred P. Kaye

## Abstract

Innate preferences determine how animals interact with the environment, but how experience refines the neural processes underlying those intrinsic motivations is not well understood. Here we develop a virtual pole descent task in which mice can repeat many trials without habituation of height-dependent avoidance. Mice adjusted their choices based on recent trial outcomes without externally imposed behavioral reinforcement or punishment. Using this paradigm, we found that noradrenergic signaling enhances height avoidance while experience refines the prefrontal cortex population representation of the task. Inhibiting locus coeruleus norepinephrine neurons reduced height avoidance to visually tall cliffs, while stimulating noradrenergic projections to prelimbic cortex enhanced safe decision-making. Anticipatory norepinephrine in prelimbic cortex correlated with height avoidance across animals and reflected trial outcome history. Miniscope calcium imaging revealed prelimbic neurons tracked progress in the task. At the population level, experience improved decoding of position from neural activity which correlated strongly with behavioral improvements. With experience, neural trajectories became less variable during the task and reliability of representations correlated with behavioral improvement. Together these results reveal that innate threat experience can induce prefrontal cortical refinement without externally imposed reinforcement. Innate behavioral preference is thus maintained while the neural processes underlying it evolve, suggesting flexibility in neural circuits for interacting with hardwired environmental motivations.

## Introduction

An animal navigating a natural environment must balance its goals with environmental risks. If in the pursuit of a goal the animal encounters a cliff, the animal must balance achieving the goal with the danger of falling. To safely navigate this challenge, the animal must refine its behavioral strategy to pursue the goal while maintaining the innate aversion to height-related danger. Understanding how neural circuits respond to environmental threats to survival may improve the understanding and treatment of anxiety-related disorders. In recent years, there has been a push to move away from common behavioral assays used to assess threat circuits due to inadequate translation between pre-clinical and clinical studies which has stalled the development of novel therapies, especially in the case of post-traumatic stress disorder^1–3^. One mechanistic reason for this mismatch is the lack of preclinical threat paradigms that can study how arousal state alters threat processing on a moment-to-moment basis.

While arousal state and neuromodulator release have repeatedly been shown to modulate sensory processing in many settings, recent tools which allow for large-scale neural recordings in freely moving behavior have allowed neuroscientists to study more complex relationships between neural computations and animal behavior. Recent work demonstrated that arousal state, as measured by pupil diameter, is able to explain large scale variations in neuronal calcium recordings^4^and that seemingly task-unrelated “noise” in neural data represents meaningful information^5^. A key regulator of arousal is the neuromodulator norepinephrine. It has been well established that norepinephrine levels increase with exposure to threats, but this perspective of arousal shaping neural activity raises the possibility that norepinephrine is not simply an indicator of the presence of a threat, but that arousal-driven modulations in norepinephrine actively shape the neural representation of threat-related stimuli.

The locus coeruleus (LC) is a small brainstem nucleus which sends extensive noradrenergic projections throughout the brain^6–11^. The LC-norepinephrine system has a well-established role in the regulation of arousal states^12–18^, modulation of sensory processing^19–24^, and processing of threats^25–29^. While long conceptualized as a global regulator of arousal and sensory gain, recent evidence including findings from our own lab^30^ suggests that norepinephrine acts in a spatially specific manner to regulate behavior ^6,7,31–34^. The prelimbic medial prefrontal cortex (PL) is densely innervated by noradrenergic LC projections^31,35–37^. Given the role of the PL in threat behavior^38–40^ and decision-making^41,42^ and the abundance of NE projections to this region, we hypothesized that changes in arousal, as measured by changes in norepinephrine release in the PL, could set the neural context in which environmental threat stimuli are encoded and influence the subsequent behavioral outcomes.

Current threat models are not suitable for answering this question. Fear conditioning paradigms result in rapid and robust learning^43^ while innate predator stimuli, such as looming, show rapid behavioral habituation^44^. As such, neither type of classic threat paradigm is well-suited to study how fluctuations in arousal alter neural responses to environmental threats which require trial-to-trial changes in arousal and the ability to experimentally repeat many trials. Another challenge of existing pre-clinical threat paradigms that may contribute to low translation is a lack of threat stimuli which evoke similar behavioral responses in both rodents and humans. For example, the primary behavioral outcome measure of fear conditioning is freezing in rodents, while outcome measures in human studies include skin conductance latency and subjective reports of fear^45^. Innate predator stimuli such as looming reliably evoke fear responses from rodents^44,46^, but human studies primarily utilize human conspecifics^47–50^ as social threats rather than ethologically-based “predator” stimuli^51^. In contrast, the fear of heights is an ethologically relevant, evolutionarily conserved visual threat that can be applied consistently in both rodents and humans.

The visual cliff task is an optical illusion which tests how subjects interact with a perceived cliff. Work using the visual cliff has shown many mammalian species, ranging from rodents to human infants^52^, avoid crossing the threshold of the visual cliff. While this assay has primarily been used to study depth perception^53–55^, this early work suggests a common aversion to height threat. More recent work which exposed humans to innate threats in virtual reality demonstrated that virtual height threat, but not virtual predator threats, induced subjective and physiological measures of arousal and anterior insula oscillations^56^. Rodent studies of height threat typically place rodents on an elevated platform and measure physiological and neural responses to being in the high place. These studies have shown that behavioral responses to height threat follow a psychometric curve^57^, height threat evokes activity of height-threat specific neurons in the basolateral amygdala and increases heart rate^58^, and defined a non-image forming visual circuit that regulates mouse behavioral responses to being on an elevated platform^59^. While informative, this paradigm is limited in the number of threat manipulations available to the researcher (changing the depth, opacity, or openness of the platform) and the behavioral responses available to the rodent (exploration of the edge, head dips, trembling or freezing) that can be studied. In humans, the use of virtual reality to study fear of heights has already been well established^60–62^. Thus, a virtual height threat paradigm for mice would be the most direct pre-clinical behavioral correlate.

To enable the study of precise threat computations and how they may evolve with experience, we built upon a recently published pole descent visual cliff task^57^. In our pole descent virtual visual cliff task, mice are placed on a pole and must descend towards a visual cliff image displayed on a computer monitor. The pole ends in an exit cone, placed on a Plexiglas surface above the monitor, which is positioned at the precipice of the visual cliff. The use of the pole descent version of the visual cliff task allows us to characterize height threat-related exit choices as safe or unsafe choices, reflecting the appropriate behavioral response were the visual cliff a true cliff. This task retains a naturalistic, ethologically relevant height threat while providing greater affordances to the mouse and our ability to interpret their behavior. In contrast with elevated platform paradigms, exposure to height threat in a virtual visual cliff paradigm can be tightly controlled by the experimenter, threat level can be parametrically controlled by changing the perceived depth, and the behavioral and neural responses to height threat do not rapidly habituate^58,59^. These features allow for rigorous characterization of neural circuits involved in the regulation of threat behavior and how they are modified by arousal.

We performed behavioral characterization and validation of the task, then used it as a tool to delineate neural mechanisms underlying behavioral control of innate threat avoidance. Behaviorally, both male and female mice avoided making height decisions with apparently high drop-offs. The repeated trial structure of this task permitted testing behavioral choice structure using predictive models (generalized linear mixed model), which identified trial-to-trial flexible adjustment of heights avoidance. Mice improve behavioral performance after prior trial unsafe exits from the pole, which has previously been linked in learning tasks to noradrenergic input to frontal cortex^63^. We combined optogenetic modulation of the locus coeruleus-norepinephrine system and its projection to prelimbic cortex, fiber photometry of norepinephrine release in prelimbic cortex, and single cell calcium imaging of prelimbic neurons to test our hypothesis that prelimbic norepinephrine dynamically regulates prelimbic control of threat avoidance behavior. We established that within our virtual pole descent task, the locus coeruleus-prelimbic circuit regulates behavioral responses to environmental threats in a manner consistent with virtual height stimuli being perceived as threatening. Additionally, we found that PL nore-pinephrine release during an anticipatory period, prior to trial initiation, reflects recent outcome safety and is predictive of behavioral outcomes early, but not late, in the experimental session, suggesting both momentary and longer-scale changes in noradrenergic control with experience. Using single cell calcium imaging, we identified single neurons in PL with dynamics reliably tied to mouse position along the pole and found that PL population dynamics became more stereotypical with experience. Improvement in mouse position decoding accuracy from neural data and enhanced neural stability in PL ensembles correlate with mouse-level improvements in threat avoidance, despite group-average threat avoidance remaining stable with experience. Together, these data demonstrate how exposure to internally driven height avoidance involves flexible adaptations in internal state and population dynamics. We demonstrate that experience refines PL population dynamics in a way that supports adaptive threat avoidance and propose that PL norepinephrine modulates the neural state in which threat stimuli are processed and acted upon.

## Results

### Validation of a Pole Descent Virtual Height Threat Task for Studying Threat Computations

We sought to establish a height threat task that was highly amenable to experimenter control. To enhance our ability to quickly and randomly present visual cliff stimuli during the pole descent task^57^, we modified the task to a virtual version (Figure 1A-C). Mice were randomly presented with low (5 cm), high (20 cm) or flat (0 cm) virtual visual cliff stimuli with both the apparent height of the stimulus and the position of the proximal quadrant randomized between trials. These visual stimuli were chosen due to clear differences in behavioral performance between them on the standard pole descent visual cliff task^57^. Thus, if the virtual height stimuli were effective, mice should choose the safe quadrant more often in the high cliff condition.

**Figure 1.**
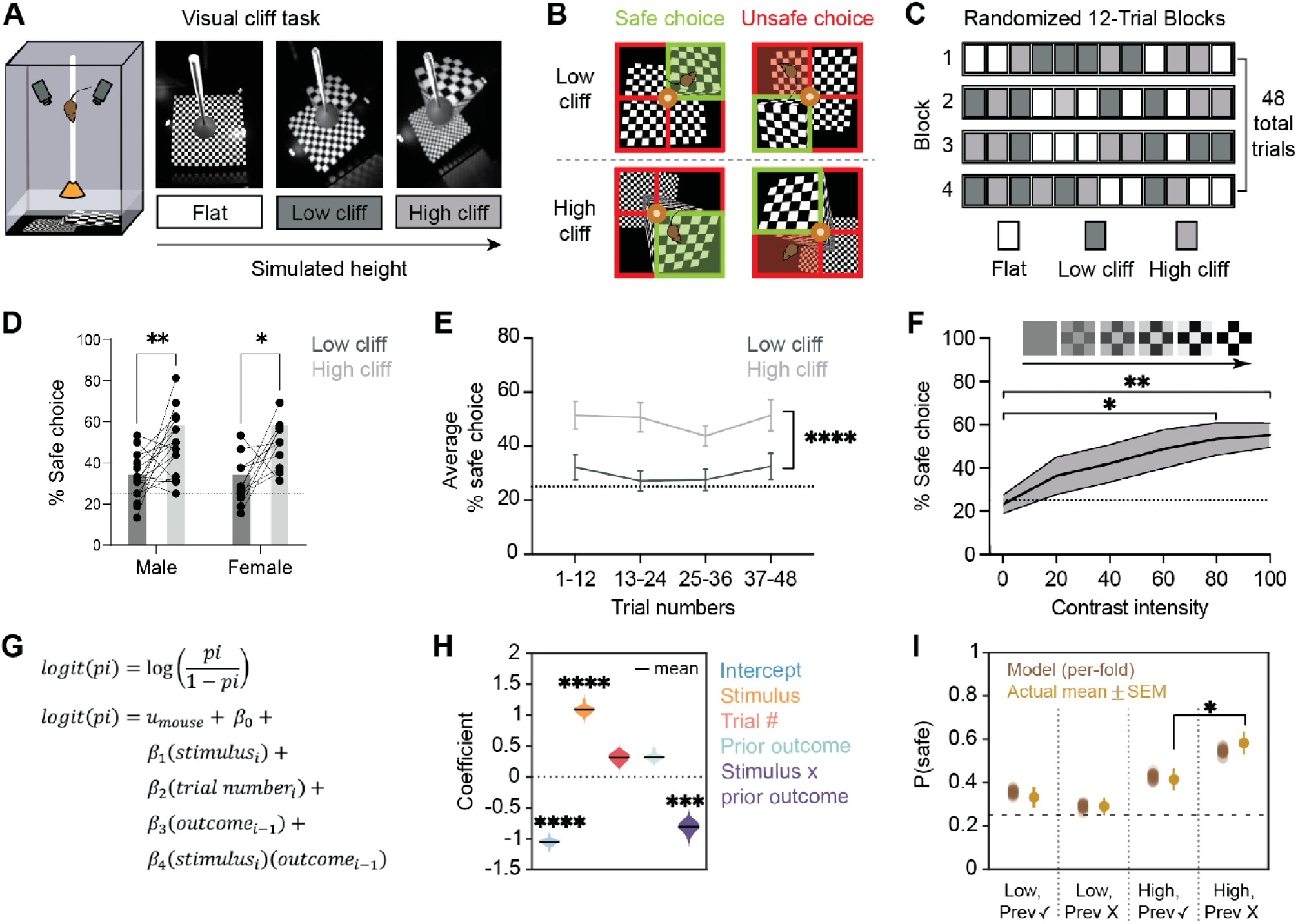
Characterization of behavior in a mouse pole descent virtual cliff task. A) Schematic of the behavioral box with example flat, low, and high cliff visual stimuli. B) Schematic describing how safe (exiting to the proximal quadrant) and unsafe (exiting to any other quadrant) outcomes are defined across visual stimuli. C) Schematic of the experimental trial structure. Stimulus height and safe quadrant are randomized in 12 trial blocks for a total of 16 presentations of each height stimulus. D) Both male and female mice make the safe exit choice significantly more often in the high cliff condition (RM Two-way ANOVA, p<0.0001 for height, sex n.s., height x sex n.s.). **E**) Safe choice behavior is consistent across all 48 trials (Two-way Repeated Measures ANOVA, block n.s., height p < 0.0001, block x height n.s.). F) Reducing the contrast of the visual stimulus reduces safe choice frequency (p = 0.05, repeated-measures ANOVA). G) A generalized linear mixed model (GLMM) with a logit link function was used to test the role of behavioral parameters on the probability of making a safe exit choice. The beta weights for current stimulus type (F(1,92) = 16.22, p = 0.0001) and interaction of current stimulus type and previous trial outcome (F(1,92) = 9.61, p = 0.003) were significant contributors to the GLMM (Type III ANOVA) (H). (I) Model predictions accurately reflect behavioral data and show increased probability in a high cliff trial if the previous outcome was incorrect (Individual repeated measures 2-way ANOVAS; current high cliff trials (F(1,228)=5.86, p = 0.02); current low cliff trials (F(1,224)=2.57, p = 0.11)).

Both male and female mice exited to the safe, visually proximal quadrant more frequently in the high cliff condition than in the low cliff condition (Figure 1D). A two-way repeated measures ANOVA showed a significant main effect of height (F(1,26) = 17.93, p < 0.0025), but not sex (F(1,26) = 0.070, p = 0.794), or a height *×* sex interaction (F(1,26) = 0.112, p = 0.740), suggesting both male and female mice discriminate between height stimuli. Post hoc Šídák tests confirmed a significant increase in exits to the safe quadrant in the high versus low cliff condition in both males (p = 0.009) and females (p = 0.0138). Given no height x sex interaction, sexes were pooled for subsequent analyses. To assess potential confounding effects of spatial preferences within the arena unrelated to the virtual visual cliff stimuli, we incorporated a flat visual stimulus with no visual depth disparity. While some individual mice displayed spatial preferences (see Methods), performance on flat stimulus trials confirms our findings were not driven by an overall spatial preference within the test arena (Supplemental Figure 1A, Repeated Measures ANOVA (F(2.641,71.31) = 1.022, p =0.38)). This finding is also true when comparing quadrant choice over all trials (Supplemental Figure 1B, Repeated Measures ANOVA (F (2.625, 70.88) = 1.596, p = 0.20)).

Although innate threats require no training to elicit a behavioral response, behavioral responses to innate threats are modified by experience and visual innate threats such as looming rapidly habituate^44,64^, limiting the ability for repeated stimulus presentations. Like a visual looming stimulus, the pole descent virtual visual cliff task is an optical illusion where each trial has a behavioral outcome free of experimenter-induced contingencies. Mice do not receive external punishment for unsafe choices or reinforcements for safe choices. Thus, it is possible mice could learn that unsafe choices have no consequences, and their threat response could habituate over the course of trials, as is observed in visual looming paradigms. We calculated the average frequency of safe exit choices for each mouse in response to the low and high cliff stimuli, separated into 12-trial blocks (Figure 1E). A two-way repeated measures ANOVA revealed a significant effect of height (F(1,22) = 23.81, p<0.001) but not block (F(3, 66) = 0.98, p = 0.41) or a height x block interaction (F(3,66) = 0.19, p = 0.901). Post hoc Šídák tests confirmed significant differences between height stimuli at each block (block 1, p = 0.01, block 2, p = 0.001, block 3, p = 0.02, block 4, p = 0.001) but no significant effect of block number on behavioral outcome within each height stimulus condition (p>0.9 for all comparisons). We found that mice maintained their behavior across the 48-trial session, consistently making safe exits more often in the high cliff condition even in the final block of the session. A subset of mice underwent a second testing day 48 hours following the first session to assess potential learning effects. We found that safe quadrant preference in the high cliff condition decreased on the second test day (Supplemental Figure 1C, Two-Way Repeated Measures ANOVA; Effect of day F(1,13) = 0.94, p = 0.38; effect of height F(1,13) = 9.93, p = 0.008; height x day interaction F(1,16) = 6.67, p = 0.02). Our data support that the virtual height threat task allows for repeated exposure to an innate threat stimulus without habituation within a session. This task uniquely allows experimenters to repeat threat exposure over many trials, facilitating robust statistical comparison of neural data. Simultaneously, this task has utility for assessing computations underlying threat learning and habituation by completing multiple behavioral sessions.

To establish that our results were due to the perception of height threat, rather than a preference for patterns with certain spatial frequencies, we modified the stimuli to have the same spatial frequency but without the illusion of height. We found that mice had no preference for the smaller spatial frequency quadrants without corresponding depth cues (Supplemental Figure 1D. paired t test, t = 1.37, df = 7, p = 0.21), supporting the conclusions of Boone et al.^57^ that depth cues are required for mice to show behavioral discrimination across visual height cues. To further verify the role of vision in task completion, we altered the contrast of the visual stimulus from 100% to 0%, hypothesizing that as the visual stimuli became harder to see, the increased frequency of safe exit choices in the far condition would decrease. Indeed, when tested with high cliff stimuli of various contrast intensities, mouse preference for the safe quadrant decreased as a function of stimulus contrast (Figure 1F). A repeated measures ANOVA revealed a main effect of contrast on behavioral outcome (F(7, 35) = 2.32, p = 0.05) with posthoc Tukey’s multiple comparisons test supporting a significant difference between 0 and 100% contrast (p < 0.001) and a difference approaching significance between 20 and 100% contrast (p = 0.06). Together, these data demonstrate mice of both sexes display stable behavioral responses to virtual height threat which is modified by experience, depth cues, and visual contrast.

We sought to determine if our pole descent virtual height threat task could be used to study the neural computations underlying threat decision-making by applying common analysis methods used in appetitive decision-making tasks. As previously mentioned, innate threats such as looming rapidly habituate over repeated presentations, limiting our ability to expose mice to many looming trials. Even in learned threat tasks, such as cued fear conditioning, mice form strong fear memories with very few threat exposures^65,66^, limiting the number of trials that can be studied to understand neural computations underlying changes in threat behaviors. Appetitive decision-making tasks, in which mice will repeat many trials for rewards, have been intensely studied with computational models to understand the processes underlying decision-making^67–69^, but few paradigms are available for studying threat-related decisions. We thus used a generalized linear mixed model (GLMM, Figure 1G) to determine what behavioral factors influenced choice behavior on the current trial. We modeled a variety of behavioral components, including the stimulus type, trial number, previous trial outcome, and the interaction of the current stimulus type and previous trial outcome. The stimulus height during the current trial significantly influences choice behavior (Figure 1H), as is seen in the averaged behavioral data (Figure 1D). Similarly, supporting our data showing choice stability over trials within a session, the trial number was not a strong predictor of current trial performance. Interestingly, the GLMM revealed that the outcome of the previous trial influenced the outcome of the current trial only during high cliff stimulus presentations (Figure 1H). A type III ANOVA conducted on the GLMM beta weights for each variable confirmed a significant main effect of current trial stimulus type (F(1,92) = 16.22, p < 0.001) and no effect of previous trial choice (F(1,92) = 1.54, p = 0.22), but a significant interaction between current stimulus type and previous trial outcome (F(1,92) = 9.61, p = 0.003). The GLMM model predictions reliably match true behavioral data (Figure 1I). Individual repeated measures 2-way ANOVAs of the GLMM beta weights supported a significant effect of previous trial outcome on current high cliff trials (F(1,228)=5.86, p = 0.02) but not for current low cliff trials (F(1,224)=2.57, p = 0.11). Our modeling results suggest that while animal performance on the task is relatively stable over 48 trials, recent unsafe outcomes are associated with increased accuracy on the subsequent trial on a trial-to-trial, stimulus specific basis, a phenomenon described in human decision-making data as early as 1966^70^. This post-error boost has been studied in appetitive tasks^71^, but to our knowledge it has not been reported in aversive tasks with similar cognitive or “decision-making” components. The noradrenergic system is one putative mechanism by which a threat-related error could rapidly boost task performance on the subsequent trial^72^.

### The Locus Coeruleus is Required for Virtual Height Threat Avoidance

Our modeling results suggest that while behavior is relatively stable when averaged across trial blocks, behavioral outcomes fluctuate on a trial-to-trial basis in a way which is driven by the interaction between a prior “error” (unsafe choice) and the current threat level. Importantly, there are no experimenter-induced consequences following the mouse’s exit that can explain this outcome-based change in task performance. One interpretation is that these outcome-based changes, occurring in the absence of experimenter-induced reinforcement, reflect an innate preference to avoid the dropoff of the visual cliff, potentially driven by the mouse’s internal drive to reduce threat-related fear. Prior literature has suggested norepinephrine could mediate post-error boosts in performance^72^ and the Yerkes-Dodson law^73^ has long described a relationship between norepinephrine levels and task performance. As a key regulator of arousal state and known modulator of threat responses, we hypothesized the locus coeruleus-norepinephrine system could regulate behavioral responses to height threat in a manner which could be rapidly modified by behavioral state.

To determine if the locus coeruleus (LC) was required in the pole descent virtual visual cliff task, we used optogenetics in DBH-Cre mice to specifically inhibit norepinephrine neurons in the LC using the inhibitory opsin stGtACR2^74^(Figure 2A). For these experiments, the cell bodies of LC noradrenergic neurons were silenced throughout the duration of laser on trials. The 48 trials were split evenly across height stimuli then separated into laser on and laser off trials to allow for within-animal comparisons. When LC norepinephrine neurons were inhibited, mice significantly reduced their preference for the safe quadrant in the high cliff condition (Figure 2D). A two-way repeated measures ANOVA revealed a significant effect of optogenetic stimulation (F(1,10) = 5.01, p = 0.05) but not of height (F(1,10) = 3.88, p = 0.08) or a stimulation x height interaction (F(1,10) = 2.008, p = 0.19). Post-hoc analyses using Fisher’s LSD revealed no effect of laser in the low cliff condition (p =0.33), but a significant effect of laser in the high cliff condition (p = 0.01) as well as a significant difference in safe choice across heights in the laser off (p = 0.001) and laser on (p = 0.04) conditions. These results confirm that LC norepinephrine neuron activity is required for normal height-threat avoidance decisions, specifically under high threat conditions. However, global somatic LC inhibition does not provide a mechanistic understanding of where and how norepinephrine is regulating height threat avoidance behavior.

**Figure 2.**
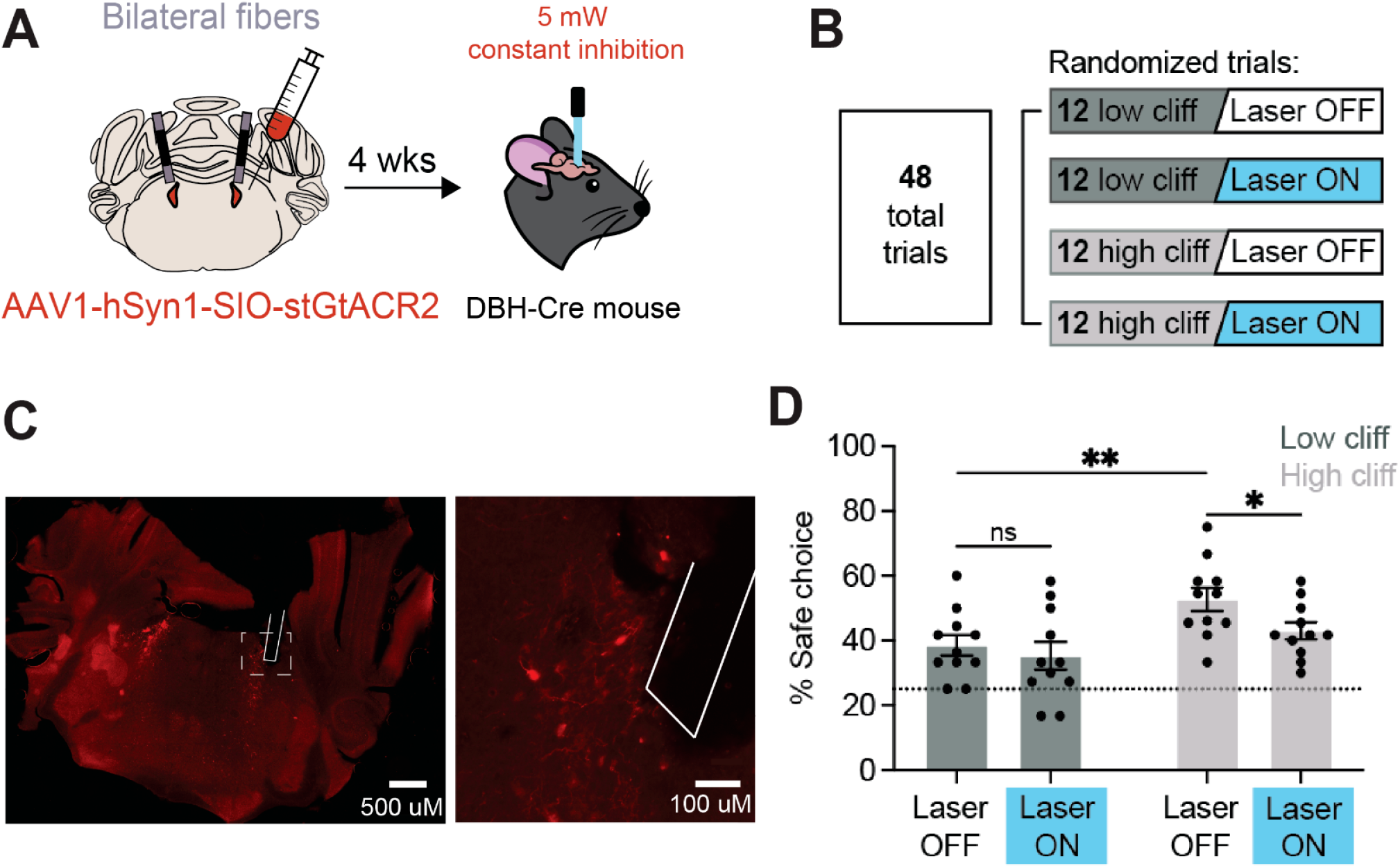
The locus coeruleus is required for safe exit choices. A) Schematic of viral injection of AAV1-syn-SIO-stGtACR2FusionRed into the LC of DBH-Cre mice for specific inhibition of noradrenergic cell bodies. B) Schematic of trial structure in optogenetics experiments. Flat trials were removed to allow for a sufficient number of Laser On/Laser Off trials for each height stimulus. C) Image showing viral expression and placement of optogenetic fibers in the LC. D) Optogenetic inhibition of LC noradrenergic neurons decreases frequency of safe choice in the high cliff, high threat condition (Main effect of stimulation (F(1,10) = 5.01, p = 0.05, repeated measures 2-way ANOVA. Posthoc effect of laser in the high stimulus condition, p = 0.01).

### A LC – PL Circuit Regulates Height Threat DecisionMaking

Optogenetic inhibition of LC norepinephrine neuron cell bodies significantly reduced safe exit choice behavior in this task. This verifies the involvement of the LC norepinephrine system in height avoidance behavior but doesn’t demonstrate how LC norepinephrine may influence downstream structures to guide behavior. The prelimbic prefrontal cortex (PL) is a key region for regulating decision-making behavior, is activated by spatial proximity to threat, and receives dense noradrenergic projections from the LC^37^. To directly test the role of the LC-PL noradrenergic projection, we optogenetically stimulated LC-NE terminals in the PL of DBH-ChR2 or control (DBH-Cre (-), flex-ChR2(+)) mice (Figure 3A). We found that stimulation of LC-NE terminals in PL enhances safe choice frequency across height conditions in DBH-Cre positive, but not DBH-Cre negative, mice (Figure 3B). A three-way ANOVA revealed a significant main effect of genotype (F(1,8) = 11.16, p =0.01) and height (F(1,8) = 8.095, p = 0.02) but not stimulation (F(1,8) = 0.58, p = 0.47). There was a significant genotype x stimulation interaction (p = 0.007), but not genotype x height x stimulation interaction (F(1,8) = 0.89, p =0.37), confirming that optogenetic stimulation of PL norepinephrine release altered behavior across height stimuli in a genotype dependent manner. Post-hoc testing with Sidak’s multiple comparisons test between genotypes confirmed a significant difference between control and DBH-Cre x flex-ChR2 mice in both low cliff (p = 0.03) and high cliff (p = 0.02) trials only when the laser was active (low cliff p = 0.97, high cliff p = 0.97 between genotypes in laser off condition).

**Figure 3.**
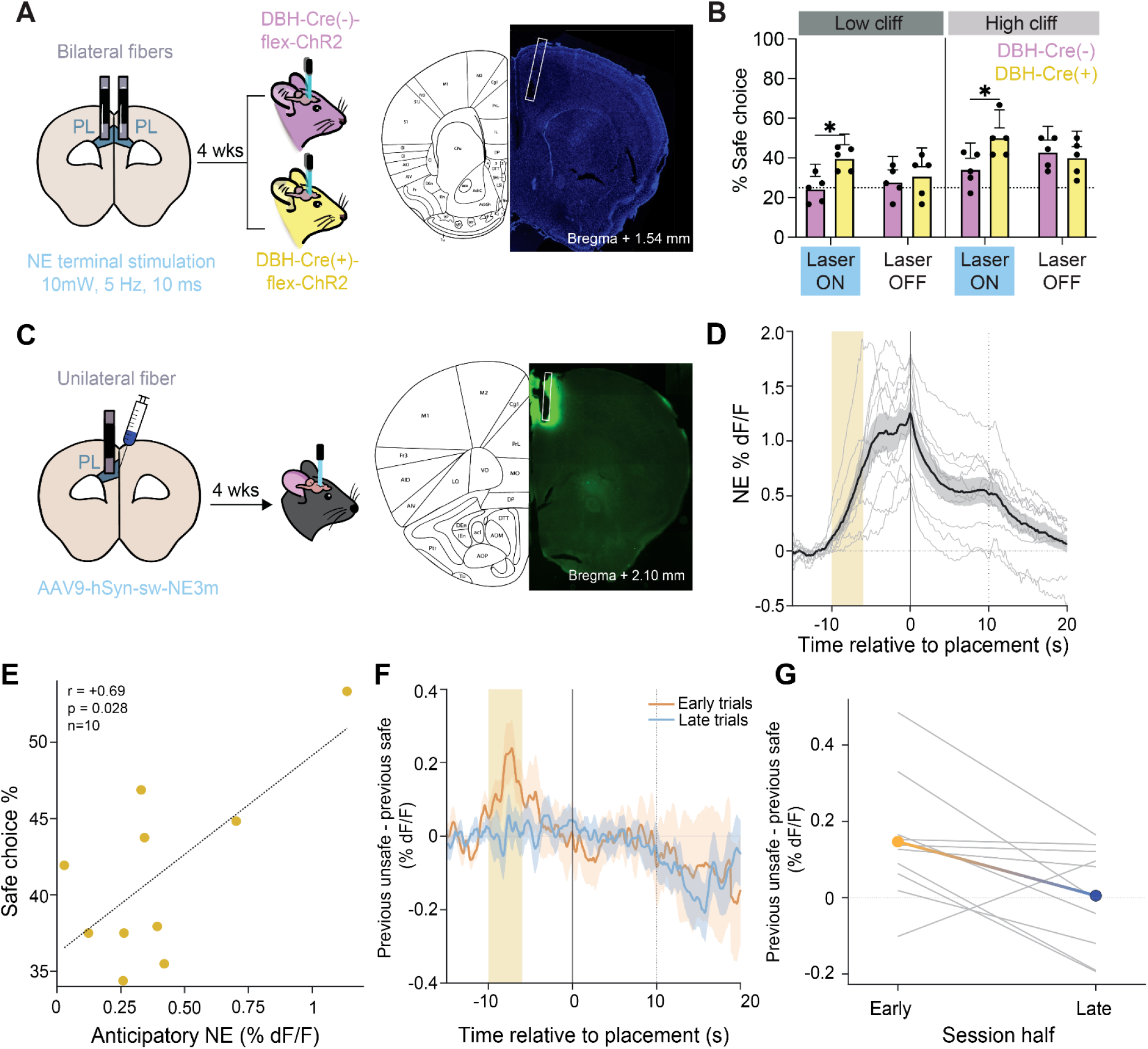
Prelimbic noradrenergic release reflects outcome safety and adapts with experience. A) Schematic of surgical and optogenetic procedures. Fibers were implanted bilaterally in the PL of flex-ChR2 mice crossed with either DBH-Cre negative or DBH-Cre positive mice to allow for specific opsin expression in DBH-expressing neurons. B) Optogenetic stimulation of LC terminals in PL increased safe choice frequency in response to both the low cliff and high cliff stimulus. Optogenetic laser had no effect on DBH-Cre negative mice which do not express ChR2 (Three-way ANOVA, genotype (F(1,8), = 11.6, p = 0.01), height (F(1,8) = 8.095, p = 0.02), stimulation n.s., genotype x stimulation (p = 0.007), genotype x stimulation x height n.s.) C) Schematic of viral injection (AAV9-hSyn-sw-NE_3m_) and fiber optic implant for fiber photometry recordings of NE release in PL. D) Mean (black) and SEM (shading) PL NE fluorescence across mice. Gray lines are individual mouse means. Mouse placement on pole is time 0 and exit is warped to +10 seconds. The anticipatory window is −10 to −6 seconds, shaded in orange. E) Correlation between mean anticipatory NE versus overall safe-choice percentage (Pearson r=0.699, two-sided p = 0.024, n=10). F) The difference in norepinephrine levels during the anticipatory period following safe versus unsafe trials changes over experimental session. Mouse means for the unsafe – safe difference in NE dF/F for early trials (orange) versus late trials (blue). G) Paired anticipatory differences for each mouse (gray) and the group mean (orange to blue gradient, session half *×* previous outcome F(1,9)=10.82, p = 0.009). Trials following flat stimuli are excluded from F–G.

### Prelimbic norepinephrine signals prior errors in an experience-dependent manner

Given that stimulating noradrenergic terminals in PL enhanced performance, we reasoned that norepinephrine dynamics in PL during the task would show reproducible task-related dynamics. The GPCR-based sensor GRABNE_3m_ was expressed in PL neurons and an optical fiber was implanted to record NE concentration dynamics in real-time during the task (Figure 3C). Naturalistic behaviors involve trial-to-trial variation in timing. Linear time-warping^75^(see Methods) was used to standardize norepinephrine fluorescence across trials to a standard duration between placement on the pole and exit. Norepinephrine concentration rose during the period before being placed on the pole and remained high throughout the trial (Figure 3D). PL norepinephrine levels then gradually decreased once the mouse exited the pole onto the plexiglass surface (Figure 3D). Thus, norepinephrine extracellular concentration in PL is elevated by the virtual height threat decision task.

Prior work in our lab has demonstrated that norepinephrine release in PL represents a threat prediction error, providing PL with information about threat expectations^76^. We observed norepinephrine release in PL increases before the mouse is placed on the pole. We defined the period of −10 to −6 seconds prior to placement as the anticipatory period, where the mouse is still in a holding cage but will soon be picked up to start a trial. The amount of norepinephrine release during this anticipatory period positively correlated with threat avoidance behavior, with mice with higher anticipatory norepinephrine having a higher percentage of safe exit choices (Figure 3E, Pearson r=0.699, p = 0.024, n=10).

Interestingly, we did not see clear differences in the amount of PL norepinephrine release with respect to current stimulus type or behavioral outcome when comparing mouse-averaged data (Supplemental Figure 2A). We hypothesized that norepinephrine dynamics would shift as mice had repeated experience in the task. To test this, we split each session at the median trial index. To test if previous errors might be reflected in PL norepinephrine release, we labeled trials by the outcome of the preceding trial. Anticipatory nore-pinephrine was higher following unsafe than safe outcomes early in the session, and this previous-outcome difference diminished in the second half (Figure 3F-G. session half *×* previous outcome, F(1,9)=10.82, p = 0.009). Thus, differences in PL norepinephrine release tracked prior outcome safety rather than predicted future outcomes. Together, these results show that apparently stable average avoidance can coexist with experience-dependent changes in the neural state accompanying threat encounters.

The heightened activity upon placement on the pole replicates previous literature demonstrating high amounts of norepinephrine release in response to tail lift and presentation of an experimenter hand^77,78^. Although the current behavioral paradigm does not allow us to dissociate the effect of experimenter handling from height exposure in terms of norepinephrine release, we performed fiber photometry of nore-pinephrine release in PL during an elevated platform task to assess the effect of height exposure itself on norepinephrine release in PL. In this task, mice can freely move on an elevated stationary disk. We analyzed changes in PL norepinephrine release as mice looked over the edge of the stationary disk and found a similar pattern of high PL norepinephrine release as mice peered over the edge of the disk, directly engaging with the visual height threat, which slowly returned to baseline levels once mice moved away from the disk’s edge (Supplementary Figure 2B). Thus, visual height stimuli evoke PL norepinephrine release independent of experimenter handling.

To determine if damage to the PL or connection to implant cables disrupted behavior, we compared the behavior of control mice, mice with an optic fiber implanted in the PL, and mice with a GRIN lens implanted in the PL. We found no differences in exit choice behavior across these surgical conditions, confirming surgical preparations did not alter task behavior (Supplemental Figure 1E; Repeated Measures 2-Way ANOVA, for height (F(1,36) = 25.06, p <0.0001), implant group (F (2,36) = 0.19, p = 0.83), and height x implant group (F (2,36) = 0.64, p = 0.53)).

### PL population activity during virtual height avoidance

PL cortex mediates choices in both appetitive and aversive tasks and may play a particular role in tasks involving flexible adjustment of conflicting motivations. Thus, we next sought to determine how PL population activity integrates task information to control virtual height threat behavior. To test this, we used single cell microendoscope calcium imaging of PL neurons during the virtual height threat task. As with photometry recordings, we applied linear time-warping (see Methods) to standardize trial lengths for visualization. On a population level, we find sustained activity of PL neurons across the time from placement to exit, relative to the pretrial baseline period (Figure 4D).

**Figure 4.**
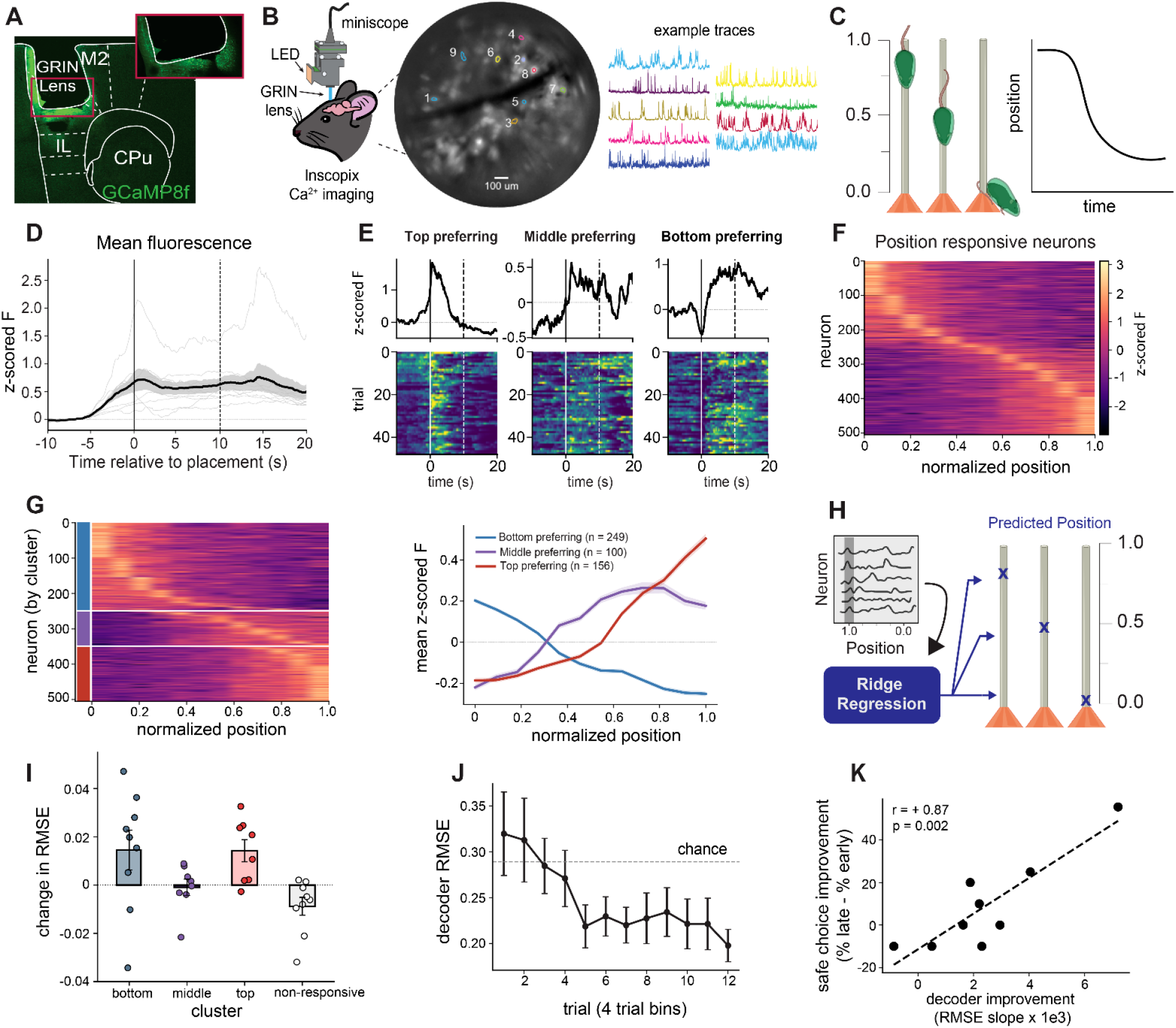
PL population activity tracks descent progression and refines with experience. A) Example histological image of GRIN lens placement and GCaMP expression in PL. B) Schematic of miniscope imaging with example ROI and individual neuron calcium traces. C) Schematic of SAM 3 computational tracking of mouse position. D) Average PL fluorescence across mice (black, SEM shading) with individual mice in gray. Mouse placement is at time 0 and exit is warped to +10 seconds. E) Exemplar top-preferring, middle-preferring, and bottom-preferring neurons, with trial-mean traces above and individual-trial heatmaps below. F) Heatmap showing z-scored fluorescence for position tuning neurons (505/985 neurons across nine mice) sorted by preferred position. Position ranges from pole bottom (0) to pole top (1). G) Position responders sorted by k-means clustering (left) and mean cluster fluorescence over position (right, *±* neuron SEM). H) Schematic showing mouse position decoding using neural data, SAM 3 positions, and ridge regression. I) Change in ridge regression RMSE after removing individual clusters shows bottom and top position clusters have highest impact on decoder accuracy (Bars are mean *±* SEM, dots represent mice). J) 4-trial bin contributions to decoder RMSE (mean *±* mouse SEM) show that decoder accuracy improves over the experimental session. K) Position decoder improvement (a negative RMSE slope) versus safe-choice improvement (percent safe exit in last minus first third of trials). Decoder improvement is positively correlated with improved threat avoidance (Pearson r=0.875, p = 0.002, n=9).

Qualitative observations of single neurons across trials (Figure 4E) suggested distinct position-based response profiles. To understand if individual PL neurons have positionrelated activity, we analyzed individual neuron responses along the mouse’s descent trajectory using computational tracking of mouse position (Segment Anything Model 3^79^, Figure 4C). Position-shift permutation tests identified 505 of 985 neurons with significant activity–position associations (Figure 4F, FDR q<0.05, see Methods). Clustering (k-means clustering, k=3) showed a continuous distribution of position representation in the neuron-level responses for the top, middle, and bottom of the pole, with greater numbers of neurons showing position-locked activity to the top and bottom of the pole compared with the middle (Figure 4G). Similar activitytime associations are observed when neural data is warped to a common time frame (0s placement on pole, 10s decision, Supplemental Figure 3), thus this neuron-level tuning does not distinguish physical location from temporal progress towards avoiding threat and leaving the pole.

To better understand the innate representation of pole descent, we used a ridge-regression model to test if we could predict continuous position from neuron level data within mice. Ridge-regression performed well at predicting position along the pole relative to other models. By computationally “ablating” selective neuronal clusters, we compared the contributions of each position-based cluster to the overall position decoding accuracy. We determined that neurons preferring placement (top) and exit (bottom) spatial positions contributed most strongly to the decoding (Fig. 4I). Interestingly, training the model only on periods when the mouse was at the top and bottom of the pole was sufficient to enable decoding of its progress during the full task, with little change in error if the middle preferring clusters were removed.

Innate behaviors, such as the visual height threat avoidance in this task, do not require training but can still adapt over the course of experience. While no overall change in performance was noted in our task over 48 trials, we reasoned that repeated experience might involve refinement in the neural strategies employed by the mouse. When analyzed over the course of trials, decoding error gradually and significantly diminished from early to late trials (Figure 4J). Since some mice improved more than others with experience, we reasoned that this improved decoding of height descent could accompany improvements in behavior. We found that ridge regression error improvement over trials correlated with behavioral improvement (increased safe choice %) over the same period at the mouse level (Figure 4K, r=0.875, p = 0.002, n=9). To better understand the decoder improvement with experience, we tested the generalization of models trained on early trials to late trials (and vice versa). Ridge-regression models trained on early trials generalized better to late trials than the reverse, supporting a possibility that neural responses in early experience were outside of the distribution of the late responses.

### Experience Stabilizes PL Population Dynamics

Prior studies have shown that neural population dynamics change over learned behavioral training by reducing variability. Innate behaviors do not require training, yet the animal pursues an internal goal and may also refine strategies to achieve it in an analogous way. We thus sought to determine whether neural trajectories during pole descent towards a visual height choice showed reduced variability in its geometry with repeated trials.

To assess this question, we visualized neural population trajectories using principal component (PC) space (Figure 5A-B, example mouse). Within the neural trajectory space, pole descent trajectories appeared more stereotyped in late vs early trials. In late trials, there was a 56.3% geometric reduction in variance in the neural trajectories as compared with early trials (variance around mean trajectory for that time-point in PC1-2 space). To better understand this phenomenon, we conducted a Procrustes analysis to map neural trajectories to a common space between mice^80^. As with the analysis in PC space, trajectory variability diminished (Figure 5E, held-out full-space variance decreased by 58.6%, p = 0.008). The trajectories also became straighter in the neural space (straightness 0.179 early, 0.251 late; Figure 5F, p = 0.012), suggesting that refinement of neural activity patterns was not solely due to reduced variability.

**Figure 5.**
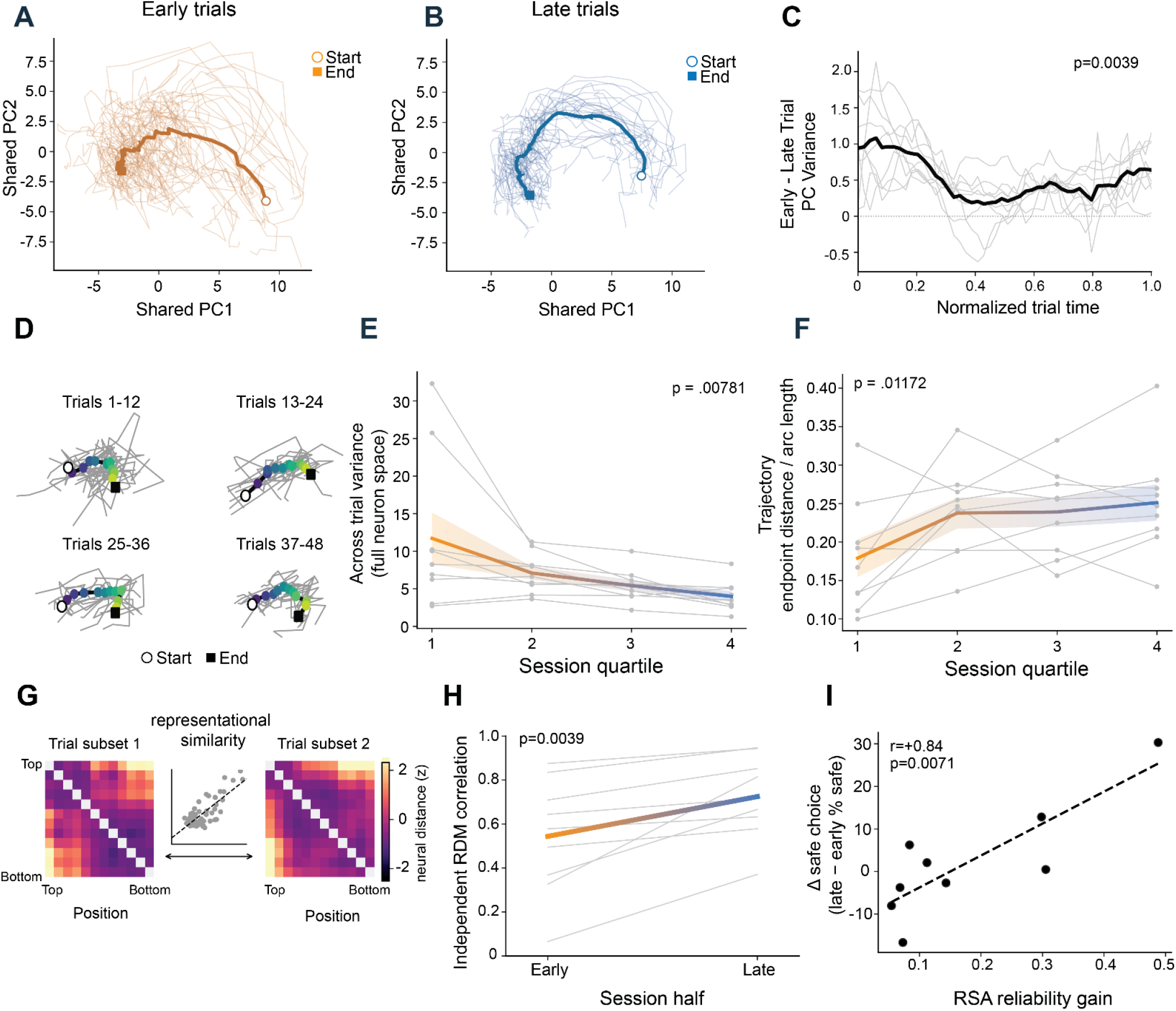
Experience stabilizes prefrontal population dynamics. A–B) Individual early (orange) and late (blue) trial trajectories in PC space for an exemplar mouse. C) Across-trial sample variance averaged across PC1 and PC2: early trials minus late trials across normalized time, divided by each mouse’s early trial mean. Reduction in the geometric mean from early to late is 56.3% (9/9 mice, exact two-sided sign-flip p = 0.004). D) All individual trial trajectories from an exemplar mouse, organized by 12 trial bins. Gray lines are trials and thick lines means, colored by position. Open circles denote the start of the trial while closed squares denote the end of the trial. E) Reduction in trial variability across full neural trajectory space (8/9 mice, exact log-ratio p = 0.008). F) Observed straightness of neural activity trajectories, defined as endpoint-to-endpoint distance/arc length (8/9 mice, p = 0.012). G) Schematic illustrating reliability analysis using two independent representational dissimilarity matrices (RDM). Each RDM combines two disjoint four-trial groups and no trials are shared between RDMs. H) Reliability between independent-RDMs increases from 0.544 to 0.725 (9/9 mice, p = 0.004). I) Improvements in RSA reliability correlate with safe-choice improvement (r=0.837, permutation p = 0.007).

We next measured how distant neural states are from one another across the pole descent neural trajectory before and after extensive height threat experience. For representational similarity analysis (RSA) two independent cross-validated distance matrices were calculated (representational dissimilarity matrix; RDM) between neural states early and late in the experimental session at each pole position. For example, for early trials, two RDMs are calculated for subsets of trials. The correlation between them serves as a metric of position-dependent neural state reliability (Figure 5G). In late trials, reliability increased in all nine mice relative to early trials (Figure 5H, p = 0.004) demonstrating strong refinement of the neural encoding of task performance. Importantly, the increased neural representation reliability correlated with improved threat avoidance (Figure 5I, r = 0.837, permutation p = 0.007). These results demonstrate that adaptive threat decisions occur in the setting of an increasingly reliable neural representation of space relevant to the innate height decision.

## Discussion

We developed and characterized a novel height threat paradigm for mice as a method for understanding momentto-moment changes in computations underlying threat behaviors. Height threat is evolutionarily conserved between rodents and humans^52,54,55,58,60^ and an ethologically relevant stimulus for both species, which is rare for threat stimuli. This trans-species behavioral component may allow for improved translation from rodent findings to humans, but also for information flow from humans to pre-clinical circuit studies. A prior study of single neurons in the mouse amygdala suggests specific neural populations responding to height threat^58^, while human work using virtual reality supports that virtual height threat, but not other virtual innate threats, is an effective threat stimulus for humans^56^. Height threat therefore is highly behaviorally relevant and may reveal evolutionary circuits responsible for avoiding danger. Circuit findings in this novel threat paradigm can be contrasted with those of fear conditioning and innate predator tasks to determine if different neural circuits are engaged by these threats and the implications for the treatment of human psychiatric disorders. Advances in freely-moving neural recording techniques have resulted in a renaissance in the study of naturalistic behaviors^81–83^, including ethological studies of mouse vision^84,85^. While naturalistic behaviors add complexity to neural recordings, they may lead to more robust, biologically fundamental findings that are better replicated across labs and species^81,86^.

Valence and arousal are conceptualized as separate dimensions of emotion. At the same time, high arousal is particularly a characteristic of anxiety and PTSD. Arousal alone can also be appetitive or neutral, serving as a cognitively enhancing state for the allocation of scarce resources. In this study, locus coeruleus NE neurons bidirectionally contributed to height-decisions. The impact of noradrenergic modulation of PL on heights decisions might be interpreted either in terms of enhancing the negative valence of the task or in allocating resources to the needed cognitive functions for making safe decisions. While mouse behavior remains stable during extended innate height exposure, the adaptations that occur in PL norepinephrine and PL neural population geometry are distinct. Anticipatory NE release to the aversive start of the trial and which is enhanced by prior unsafe choices diminishes, while PL representations become more reliable. These findings support a framework in which experience alters both noradrenergic arousal signaling and cortical neural dynamics, which may contribute to flexible threat responses. Since the task involves both affective and cognitive dimensions, these observations may reflect a transition between distinct forms of height-based decisions.

Changes in behavior and neural functions following errors are well demonstrated across a variety of behavioral tasks in both humans^87^ and rodents^88,89^. Studies ranging from cognition to motor to appetitive learning have measured behavioral and brain changes in response to errors and have demonstrated a role of mPFC^71,90,91^ in coordinating behavioral response to errors. While this literature has informed how we can optimize performance in appetitive settings, it has not addressed mechanisms controlling decisions in response to danger. Punishment is often used in these paradigms, such as reward omission or air puff, are aversive but not dangerous, making them distinct from threats where errors could be fatal. Similarly, while aversive outcomes in these tasks are experimentally induced, the type of behavioral modulation we observe in our fear of heights task is initiated from *internal* perceptions of danger. The ability to study internal perceptions of danger is particularly useful for comparison with human populations, who reliably report physiological sensations of fear when exposed to threats in virtual or augmented reality despite awareness that the threats aren’t real^56,92^. The physiological states experienced in humans exposed to virtual heights may be compared with the neuromodulatory dynamics observed in this study, in which norepinephrine increased after errors early in the task. This task thus has significant additional value in the study of fear, where classical learning paradigms (such as auditory fear conditioning) involve physical sensations of pain rather than perception of potentially dangerous environmental structure. Other behavioral models such as looming stimulus exposure can be maintained for relatively few trials without strong habituation in motivated escape behaviors, limiting the understanding of naturalistic decision-making over time. Additional studies probing the representation of height threat combined with circuit neuroscience studies could provide mechanistic insight into a fear pathway regulated by intrinsic, rather than extrinsic, detection of errors.

Recent literature has demonstrated modularity in the organization and function of norepinephrine release in the brain^6,7^. Rather than simply serving as a global arousal signal, NE release seems to have projection-specific effects on neural functioning and behavior. For example, in mice that learned a go no-go task, neurons in the LC responded to reward in different patterns across neurons, while LC neuron responses to air puff were consistent across neurons^93^. When looking at noradrenergic projections to the dorsal mPFC and motor cortex during this task, Breton-Provencher et al. found differential activation of LC NE axons with motor cortex projections being more active during lever pressing and dorsal mPFC projections being more active in response to aversive air puffs. Our study is consistent with modular functions of NE in that we show activation of NE release in prelimbic mPFC improves height threat avoidance behavior. Direct tests of how PL NE release changes neuronal reliability and relates to arousal are needed, while future studies can address the role of NE during height threat avoidance in other threat processing regions.

Neural activity in PL showed a strong relationship with progression through the task and shifted towards greater reliability with experience. The task involves multiple sensory, motor, and cognitive variables and neurons in prefrontal cortex show mixed-selectivity for different external variables^94^. Over the course of the experiment, the population-level decoding of progress along the pole (and through time) improved as the neural responses became more stereotyped. These population-level measures of neural representation of heights decision-making correlated at the subject level with more accurate (safe) choice behavior. These refinements of the PL population occurred over the course of experience and correlated with improvements in behavior over the experiment. Thus, the allocation of PL population activity to the performance of the task may show changes, here connected with innate experience, which have been linked to learning in associative tasks^95^.

The distinct forms of adaptation of PL neuronal activity and neuromodulatory (noradrenergic) input thus suggest complementary roles in repeated innate experience. Anticipatory NE reflects recent outcome safety early in the task when arousal states are likely high, whereas PL population activity becomes more stable across repeated encounters. The latter may reflect an increased signal-to-noise ratio or reduction in top-down activity as the task becomes more predictable^96^. The distinct forms of adaptation occurring in PL norepinephrine input and PL population activity suggest that the improved task representation does not derive directly from changes in the neuromodulatory input as the experiment is repeated. Rather, the NE signal seems to relate to overall task performance, while PL ensemble changes in reliability relate to task improvement. Whether this neuromodulatory input itself is causally related to the population geometric changes is an intriguing open question which may have implications for how population encoding evolves.

While LC projections are classically known to release norepinephrine, it has also been demonstrated that LC neurons release dopamine^97,98^, including in the mPFC^99^. In addition to norepinephrine, dopamine and acetylcholine are also important in signaling features about stimulus salience and novelty, as well as in regulating attention. A prior study in our lab demonstrated differential roles for dopamine and norepinephrine in the PL during an approach-avoidance looming predator stimulus task^100^. The present study focused on the role of norepinephrine due to the known role of norepinephrine in threat behaviors. Future studies could explore continuous recording strategies by extending this task over time, by using distinct stimulus manipulations, or by using electrophysiological approaches for measuring neural activity. The ability to repeatedly and controllably assay innate choices may lay the groundwork for understanding how neural representations evolve from intrinsic motivations such as the avoidance of heights.

## Acknowledgements

The authors would like to acknowledge Dr. Michael McClurkin for helpful intellectual discussion related to the manuscript. The authors would like to acknowledge Dr. Rosy Hosking and the Life Science Editors Foundation JEDI program for providing valuable editorial feedback on this manuscript and the Yale Wu Tsai Institute Imaging Core for providing access to confocal microscopy used in this manuscript.

## Methods

### Animals

78 C57BL/6J mice (42 male, 36 female, ordered directly from Jackson Laboratories), 29 DBH-Cre mice (21 male, 8 female, bred in-house using Jackson Laboratories Strain #033951 crossed with C57BL/6J), and 10 DBH-Cre x ChR2 mice (6 male, 4 female, bred in-house using Jackson Laboratories Strain #033951 crossed with Jackson Laboratories Strain #012569) were used in this study. Mice were an average of 4-5 months old at the time of behavioral experiments (mean 4.85, median 4.43). Mice with off-target anatomical placement or mice that failed to meet criterion in behavioral task (described below) were excluded from analyses. All animal studies were conducted using methods approved by Yale University’s Institutional Care and Use Committee.

### Surgical Procedures

#### General Surgical Procedures

Surgical procedures were conducted at 4-6 weeks of age. Mice were anesthetized using isoflurane anesthesia administered via a SomnoFlo vaporizer (5% induction, 2% maintenance). Animals were secured in a Kopf stereotaxic and administered carprofen (Covetrus, 5.0 mg/kg) prior to any surgical procedures. Eyes were protected with ophthalmic ointment (Systane night gel) and fur was removed from the scalp using depilatory cream (Veet). The scalp was cleaned using three subsequent scrubs of Betadine and 70% alcohol.

#### Viral Injections

For all injections and implants, craniotomies were performed over the site of injection using a micromotor drill (Foredom). Viruses were delivered using a Nanoject iii (Drummond Scientific) at a volume of 500 nL and a rate of 23 nL/second unless otherwise noted. For NE recordings, AAV9-hSyn-sw-NE_3m_ (AAVnergene) was injected into prelimbic cortex (AP +1.6, ML 0.3 DV −1.3 from the brain’s surface, relative to Bregma). For locus coeruleus optogenetic inhibition, AAV1-hSyn1-SIO-stGtACR2-FusionRed (Addgene) was injected into DBH-Cre (+) mice. Virus was injected at a volume of 100 nL at multiple sites per mouse to ensure coverage of the locus coeruleus and minimize failure of injection (AP −0.9, ML 0.9, DV −3.3 from the brain’s surface; AP −0.9, ML 0.9, DV −3.1 from the brain’s surface; AP −1.0, ML 0.9, DV −3.3 from the brain’s surface; AP −1.0, ML 0.9, DV −3.1 from the brain’s surface; all relative to Lambda). For prelimbic cortex calcium imaging, AAV1-Syn-GCaMP8f (Addgene) was injected into the prelimbic cortex as described for NE recordings. For viral surgeries in which implants did not occur simultaneously, the scalp was sutured closed.

#### Fiber Photometry and Optogenetics

For fiber photometry recordings, a fiber optic cannula (200 uM core, 3.5 mm length for PL, 4.5 mm length of LC, Neurophotometrics) was stereotaxically implanted using the coordinates described for viral injections^76^. For PL, implantation was simultaneous with viral injections. For LC optogenetics, implantation occurred 4 weeks following virus injection. The cannula was lowered using a motorized stereotaxic attachment (MDS-1, Narishige) at a rate of 100 uM/second until reaching a depth of 2.7mm from the brain’s surface, at which point the rate was reduced to 20 uM/second. Fluorescence was visualized during cannula placement to improve placement accuracy. For noradrenergic terminal stimulation, fibers were implanted bilaterally in the PL of DBHflex-ChR2 mice (AP +1.6, ML 0.4, DV −1.4 from the brain’s surface, angled 8°, relative to Bregma). The skull was scored prior to implant placement and implants were secured using Metabond dental cement (Parkell).

#### Microendoscope Imaging

Two weeks following viral injection, the skull was scored and the craniotomy over the viral injection site was repeated. A GRIN lens (4.0 x 0.5 mm, Inscopix) was stereotaxically implanted in PL (AP +1.6, ML 0.3, DV −1.3 from the brain’s surface, relative to Bregma). GRIN lenses were secured to the skull using Metabond dental cement (Parkell). Custom metal head plates were then attached to the skull using Metabond to allow for brief fixation prior to experiments for microscope attachment. Mice were administered carprofen (5.0 mg/kg) and dexamethasone (0.2 mg/kg) for two days following lens implant to reduce inflammation. Four weeks following lens implantation, mice were fitted with a baseplate (Inscopix) to allow for microscope attachment. The imaging window was visualized during baseplate attachment and placement was guided by visualization of fluorescence, cells, and vasculature. The baseplate was attached to the skull using Metabond. Once the metabond had cured, the entire headcap was covered with black dental cement (Lang Dental) to prevent light leak.

#### Behavioral Procedures

Animals were habituated to handling prior to all behavioral experiments with at least 2 days of 5-minute handling sessions. For microendoscope imaging, following each handling session, animals were briefly head fixed, fitted with a dummy microendoscope, then placed in a clean holding cage for 5 minutes for acclimation to the scope and brief head fixation. An Inscopix commutator was mounted above the behavioral arena to prevent tangling of the Inscopix microendoscope during recordings. For fiber photometry and optogenetic experiments, the patch cord was fed through the same commutator mount (with the commutator inactive) to minimize tangling.

#### Pole Descent Virtual Visual Cliff Task

The pole descent virtual visual cliff task was adapted from Boone et al. 2021^57^. A plastic dowel (2 cm diameter, 20 cm length, Grainger) was coated with silicone (FlexSeal) to allow for traction during climbing and placed within a custom 3D printed conical base. The virtual visual cliff stimuli consisted of checkerboard patterns where one quadrant appears proximal to the mouse while the remaining three quadrants appear more distant. Virtual visual cliff stimuli were presented on a computer monitor 10 cm beneath a 2 mm thick Plexiglas sheet. The pole and cone were placed at the center of the intersection of the four quadrants of the virtual visual cliff. Virtual visual cliff stimuli were created using p5.js. Low cliff stimuli had an apparent visual depth discrepancy of 5 cm between proximal and distant targets, high cliff stimuli had a 20 cm apparent visual depth discrepancy, and flat stimuli had no visual depth discrepancy. These parameters were chosen based on a combination of factors: behavioral results in Boone et al. 2021^57^, preliminary studies in our laboratories which confirmed behavioral differences between the two stimuli, and physical constraints of displaying more extreme visual depth cues within a confined monitor space. For the standard behavioral protocol, animals received 48 trials separated into 12 trial blocks, were given a one-minute intertrial rest period between trials, and a five minute rest period in a holding cage between blocks. 16 flat, 16 low cliff, and 16 high cliff stimuli were presented randomly, with the spatial position of the proximal or “target” quadrant being randomized between trials and the caveat that the first two trials for every animal were flat stimuli to allow for brief behavioral acclimation to the task. Safe exit choices are those in which mice exit the cone to the proximal, target quadrant. All other exit choices, where the mouse exits over the apparent cliff, are considered unsafe. For optogenetics experiments, flat trials were removed except for the two initial acclimation flat trials to allow for a sufficient number of behavioral trials for within-subject analyses. Animals were presented with 24 low cliff and 24 high cliff trials, split between laser on and laser off trials. Cliff distance and stimulation were pseudo-randomized such that laser stimulation did not occur more than two trials in a row. For visual contrast experiments, MATLAB was used to change the contrast of the virtual visual cliff images to increasing shades of gray to generate images ranging from 0% to 100% contrast. To allow for sufficient trial numbers, only high cliff stimuli were presented in the visual contrast experiments. Each target quadrant was presented at each contrast intensity (0, 20, 40, 60, 80, 100) 8 times randomly throughout the experiment. For spatial frequency control experiments, low cliff and high cliff stimuli were edited such that the safe and unsafe quadrants had identical spatial frequency to the original stimuli, but depth cues were removed. Exit choice was scored during the behavioral experiment. All behavioral data was verified by a secondary observer blinded to experimental conditions using behavioral video. Animals were excluded from analysis if: 1) they showed clear preference for one quadrant of the box, defined as choosing that quadrant on greater than 60% of the trials in a single block or 40% of total trials; 2) if the animal did not exit the pole after 2 minutes, the trial was ended and scored as a “timeout.” Experiments ended if animals had 6 timeout trials within a block and were excluded from analysis. Analysis and plotting of basic behavioral data were performed in GraphPad Prism Software.

#### Elevated Platform Task

The elevated platform task consists of a custom 3D printed (Prusa) PLA circle (15.24 cm diameter, .3175 cm thickness) attached to PVC pipe (1.9 cm inner diameter, 92 cm long) using cyanoacrylate glue. A three-way PVC connector was used to attach the main PVC pipe to a PVC pipe base which consisted of two “L” shaped legs. The platform was checked with a level and cleaned with disinfectant prior to every experimental session. An opaque plastic sheet was attached to the platform with clips prior to each trial to obscure the height threat and allow for baseline physiological recordings. A behavioral camera (Microsoft 360 LifeCam) was attached to a camera mount directly above the platform to allow for behavioral recording. Animals were placed on the platform with the opaque barrier in place for 5 minutes. Following 5 minutes, the barrier was removed, and animals were left to explore the platform for 10 minutes. The number of times animals performed a peer over the edge of the platform was scored from behavioral video by trained observers. A peer was defined as the mouse looking directly down from the edge of the platform such that the nose is no longer visible, and the backs of the ears were visible.

#### Generalized Linear Mixed Model

A Generalized Linear Mixed-Effects Model (GLMM) was used to predict task performance (safe vs. unsafe decisions) as a function of trial-level predictors. Models were fit using MATLAB’s *fitglme* function with a logit link. Fixed effects inputs to the model were binarized trial-related variables: low cliff or high cliff stimulus, previous outcome, and the interaction between these two variables, for each trial number *i* (within session). Trial number was normalized between 0 and 1 to allow for ease of interpretation alongside binary factors. A random intercept *u*_*mouse*_ was included to capture intersubject variability. We used leave-one-out cross validation: for each fold, we held out one mouse and fit the GLMM on the remaining mice. P values reported were averaged per coefficient across folds.

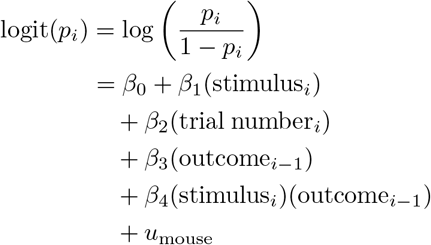

### Physiology Data Collection and Analysis

#### Fiber Photometry

Data were obtained using a FP3002 photometry system (Neurophotometrics, Inc) equipped with a 450 nM laser, 470 and 405 nM LEDs, and branching patch cord (Doric). PL NE recordings were collected at a frame rate of 40 Hz (20 Hz per channel) with interleaved excitation (470 nM) and isosbestic (405 nM) LEDs to allow for correction of movement artifacts. The LED power at the fiber tip was 100 uW^76^.

#### Fiber Photometry Analysis

A custom MATLAB script was used to align photometry and behavioral video timestamps. The time of pole placement and pole exit were scored by trained human observers. Due to the extended length of the behavioral task, trials were recorded independently and the first 10 samples of each channel (0.5 seconds at 20 Hz) were removed from analysis due to rapid photobleaching during this period. Trials were analyzed independently and first were corrected with a fourth-order, 3-Hz low-pass filter. Photobleaching was removed by fitting an exponential decay to each channel separately using robust regression with a least-absolute-residual criterion and dividing by the channel’s own fit^101^. Following photobleaching detrending, the bleach corrected signal was decorrelated from the bleach corrected isosbestic using ordinary least squares regression. To compare trials of different lengths, data in the placement to exit window were time-warped to 10 seconds using linear interpolation and a 250-ms moving average was used to smooth the interpolated trace. To allow for trial-to-trial comparisons of event-locked fluorescence, traces were baselined by subtracting the mean signal from −15 to −10 seconds prior to pole placement from the decorrelated and detrended signal. The subsequent −10 to −6 second window was defined as the “anticipatory period” in which mice had not yet been placed on the pole. Approximately 5 seconds prior to placement, mice were picked up from a holding cage and placed onto the pole.

#### Anticipatory Norepinephrine vs Behavioral Performance

The relationship between anticipatory norepinephrine release and overall safe exit percentage was quantified by Pearson’s correlation coefficient across mice. The reported p-value is a two-sided Pearson test.

#### Effect of trial history

To test the effect of prior outcomes on norepinephrine dynamics, each trial was labeled by the safety outcome of the preceding trial. Trials that were preceded by a “Flat” stimulus trial were excluded from this analysis. The remaining trials were split at the median trial index into early and late halves. The mean anticipatory norepinephrine fluorescence was calculated for each trial and a difference score was calculated for prior-unsafe trials minus prior-safe trials, resulting in one difference score per mouse per half. Changes in anticipatory norepinephrine were tested with a 2 *×* 2 repeated-measures ANOVA with session half (early, late) and prior outcome (unsafe, safe) as within-session factors.

#### Optogenetics

Trials were separated into stimulation off and stimulation on trials and stimulation was pseudorandomized such that stimulation didn’t occur for more than two sequential trials. For stimulation on trials, stimulation occurred immediately prior to being placed on the pole and continued until the decision point when the mouse exited to the Plexiglas. Animals received bilateral stimulation via the 450 nM laser (LC somatic inhibition constant illumination, 5 mW power at the fiber tip^74^; LC terminal activation 5 Hz, 10 ms window, 10 mW power at the fiber tip^102^).

#### Microendoscope Imaging

Imaging was performed using an Inscopix nVoke microendoscope connected to a commutator (Inscopix). All animals were habituated to handling and wearing the miniscope prior to behavioral experiments. For each behavioral session, a 5-minute baseline recording was performed in a holding cage prior to experiment start. For each trial, the recording started 10 seconds prior to the mouse being placed on the pole and continued for 10 seconds following the mouse’s exit. Gain and exposure parameters were chosen to minimize LED power. Videos were collected at 30 Hz and spatially (by a factor of 2) and temporally (by a factor of 3) downsampled for analysis. Preprocessing including spatial band pass filtering, motion correction^103^, and cell detection via CNMFE^104^were performed using Inscopix Data Processing Software. Identified putative neurons were manually verified based on spatial and functional characteristics. Post-processing was conducted on deconvolved df/noise traces. Downstream analysis was completed using custom MATLAB and Python scripts.

#### Time Warping

For each mouse, fluorescence traces were aligned to two main behavioral events: placement on the pole and exit from the cone. Because the interval between placement and exit varied in duration across trials, the fluorescence traces were time-warped to a fixed interval of 10 seconds based on the median trial length for this cohort (mean 11.98 seconds, median 9.93 seconds, SD 3.41 seconds across mice). The 10 seconds preceding placement and the 10 seconds following exit were not warped, resulting in 30 second-long aligned trials. The calcium imaging cohort contained ten mice and 1,084 neurons. One mouse was excluded from position analyses due to behavioral video quality, resulting in nine mice and 985 neurons in the position analyses. For the population mean, a z-score was calculated for each neuron using its mean and standard deviation over −10 to −5 seconds prior to placement, pooled across trials. Neurons were then averaged across trials, trials were averaged within mice, and finally a population average was calculated across mice.

#### Position Tracking

Each mouse’s position as it descended the pole was determined using computational behavioral analysis with Segment Anything Model 3 (SAM3)^79^. The top of the pole, bottom of the pole, and mouse centroid were marked for a single frame for every trial. A custom Jupyter notebook pipeline allowed for batch frame labeling and computational position tracking. Outputs of SAM 3 included a 1 to 0 index of mouse centroid position (top to bottom of pole) as well as behavioral videos with SAM 3 mask overlaid for visual inspection. All tracking videos were visually inspected by a human observer prior to being used for downstream analyses.

#### Neuronal Position Responsivity Analysis

We tested associations between PL neuron activity and normalized position along the pole to determine if PL neurons had specific positional responses. Fluorescence for each cell was correlated (Spearman) with position. Position selective neurons were defined as those with significant BenjaminiHochberg correction (q<0.05) relative to 1000 reshuffled circularly permuted trial shifts. K-means clustering (k=3, 20 initializations) was used to define clusters with minimum error for the n=505 selective neurons.

#### Ridge regression decoding

A ridge regression decoder with penalty *λ*=1 was trained using unwarped neural fluorescence to predict continuous position along the pole for each trial for each mouse. Mouse position ranges from 0 (bottom of pole) to 1(top of pole). The regression model was fit using 5-fold cross-validation and quantified as decoded position root-mean-square error (RMSE) on the held-out trials. For cluster removal analysis, the ridge model was refitted after removal of each cluster and RMSE was assessed on the held-out trials.

#### Population Geometry Analysis

PCA analysis: For Figure 5A–C, neuron-level fluorescence was mean-subtracted and scaled by root-mean-squared variance. Then, 50 time points within each trial were used to calculate PCA jointly on all trials as well as separately on first and second half trials. For Panel C, variance was calculated in PC1 and PC2 for early and late trials at the mouse level and then a difference score was calculated between early and late trials. For Figure 5D (visualization), individual position-binned trajectories are shown for one mouse in PC1-PC2 space for quartiles of trials. For full neural space analysis (Figure 5E–F), analysis was conducted in the full-dimensional neuron space. For panel E, sample variance calculated across held-out trials was averaged over neurons and positions within each quartile. Observed straightness was quantified as the endpoint distance divided by the sum of successive population-vector distances in full neural space, retaining the separately established analysis, scaling and trial weighting. For representational similarity analysis, a representational dissimilarity matrix (RDM) was constructed by calculating Euclidean (L2) distances between the population responses at 12 positions in full neural space. One third of trials were reserved for training and the remainder were divided into 4 groups (2 early, 2 late) for testing. The Pearson correlation between the distances of each pair (2 early or 2 late) was used to define reliability (Figure 5G) within early or late trials.

### Histology

#### Anatomical Verification

Animals were deeply anesthetized using isoflurane and underwent cardiac perfusion (10% formalin). The head, including the implant, was then post-fixed in 10% formalin for 24 hours to enhance visibility of implant tracts. Following removal, brains were sliced in 60-80 uM thick slices using a vibratome (mPFC sections, Leica) or cryoprotected in 30% sucrose and sliced in 60-80 uM thick slices on a cryostat (LC sections, Leica). Tissue was mounted directly to slides and coverslipped using Vectashield mounting media with DAPI (Vector labs). Anatomical placement was verified using an ECHO Revolution light microscope or Zeiss confocal microscope.

#### Immunohistochemistry

Animals were deeply anesthetized using isoflurane and underwent cardiac perfusion (4% paraformaldehyde). Brains were placed in paraformaldehyde overnight then moved to a 30% sucrose cryoprotectant solution. 40-60 uM thick slices were collected using a cryostat (Leica) and slices were placed into 12 well plates containing immunohistochemistry baskets and PBS for free-floating IHC (PL slices) or directly mounted to Superfrost Plus slides (ThermoFisher) for slide-mounted IHC (LC slices).

Free-floating IHC was performed for PL slices from DBH-Chr2 mice on a shaker at 70 rpm at room temperature in the dark. Tissue was washed in PBS (3, 5-minute washes) then incubated in a blocking solution (0.1% Triton X-100, 3% Bovine Serum Albumin, PBS, ThermoFisher) for two hours. Tissue was incubated in blocking solution and primary antibody overnight (1:1000 chicken anti-TH, Aves Labs, 1:2000 rabbit anti-GFP Alexa Fluor 488). The following day, tissue was washed (3, 5 minute washes) in blocking solution then incubated with a secondary antibody (1:500 goat antichicken Alexa Fluor 594, ThermoFisher; ThermoFisher) for 3 hours. Tissue was washed (3, 5 minute washes) in PBS then mounted to slides, dried, and coverslipped using Vectashield (Vector Laboratories) mounting media with DAPI.

Slide-mounted IHC was used for LC slices to preserve slice integrity. Slides were stored in a dark humidity chamber (Thomas Scientific) for the duration of the protocol. Individual slices were contained in wells using a hydrophobic pen (Vector Laboratories). Slices were washed in PBS (3, 5 minute washes) then covered with a blocking solution for one hour at room temperature (DBH-ChR2: 3% Bovine Serum Albumin, 5% Normal Goat Serum, 0.03% Triton X-100 in PBS; LC stGtACR2: 3% Bovine Serum Albumin, 3% Normal Goat Serum, 3% Normal Donkey Serum, 0.03% Triton X-100 in PBS). Blocking buffer was then replaced with primary antibodies (DBH-ChR2: 1:1000 chicken anti-TH in blocking buffer, 1:2000 rabbit anti-GFP Alexa Fluor 488; LC stGtACR2: 1:1000 chicken anti-TH, 1:1000 rabbit anti-TagRFP in blocking buffer) and stored at 4 C overnight. The following day, tissue slices were washed with 3, 5-minute washes of PBS. Secondary antibody incubation (DBH-ChR2: 1:500 goat anti-chicken Alexa 594 in blocking buffer without Triton X-100; LC stGtACR2: 1:500 goat anti-chicken Alexa Fluor 488, 1:1000 donkey anti-rabbit Alexa Fluor 647 in blocking buffer without Triton X-100) was performed for 2 hours at room temperature. Tissue sections were washed again with 3, 5-minute PBS washes prior to coverslipping with Vectashield with DAPI. IHC images were collected via ECHO Revolution light microscope or Olympus FV3000 confocal microscope.

---

**Supplemental Figure 1.**
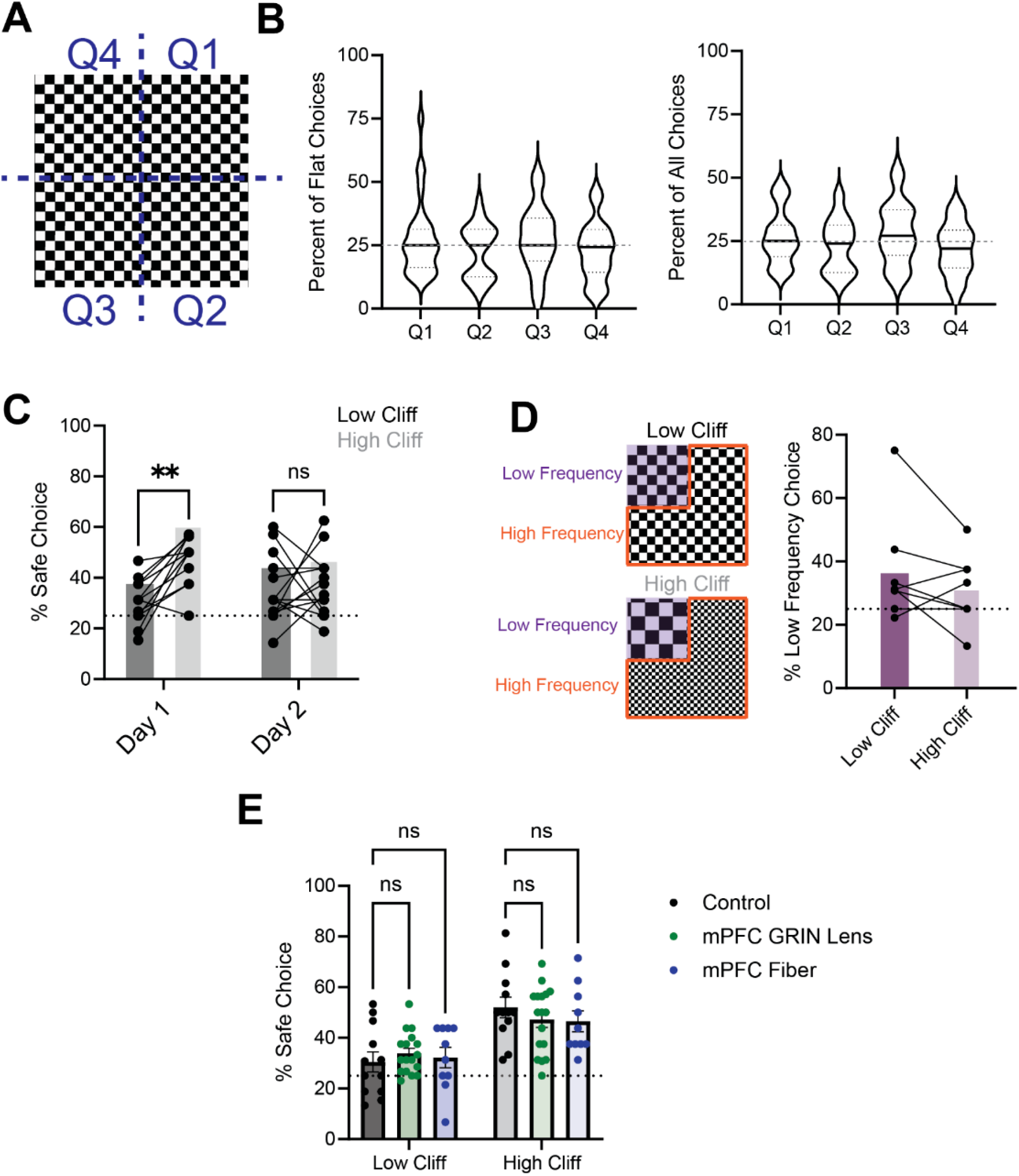
Extended Quantification of Virtual Visual Cliff Task Behavior. A) Schematic of how quadrant numbers were identified. B) Violin plot showing mean, quartiles, and distribution of mouse quadrant choice on flat trials (left). Repeated measures ANOVA demonstrated no effect of spatial quadrant on choice in the flat stimulus condition (F(2.641,71.31) = 1.022, p =0.38). Violin plot showing mean, quartiles, and distribution of quadrant choice across all visual stimuli (right). Repeated measures ANOVA demonstrated no effect of spatial quadrant on exit choice when combining quadrant choice across all stimuli (Repeated Measures ANOVA (F (2.625, 70.88) = 1.596, p = 0.20). C) Repeated experience of the virtual visual cliff task on a second experimental day reduced safe quadrant preference in the far stimulus condition (Two-Way Repeated Measures ANOVA, effect of day F(1,13) = 0.94, p = 0.38; effect of height F(1,13) = 9.93, p = 0.008; height x day interaction F(1,16) = 6.67, p = 0.02). D) To determine that mouse behavior was due to the perception of height rather than a preference for spatial frequency, we removed the depth cues from the visual stimuli and tested for preference of the low frequency quadrant. We found no difference in preference for the low frequency quadrant between conditions, confirming depth cues were necessary for preference of the low spatial frequency quadrant (paired t test, t = 1.37, df = 7, p = 0.21). To determine that implanted hardware or connection to a tether did not alter choice behavior, we compared the behavior of control mice with no implant or tether to that of mice implanted with a GRIN lens or optic fiber in the PL mPFC. We found physiology implants did not affect behavior in the virtual visual cliff task. (Repeated Measures 2-Way ANOVA, for height (F(1,36) = 25.06, p <0.0001), implant group (F (2,36) = 0.19, p = 0.83), and height x implant group (F (2,36) = 0.64, p = 0.53).

**Supplemental Figure 2.**
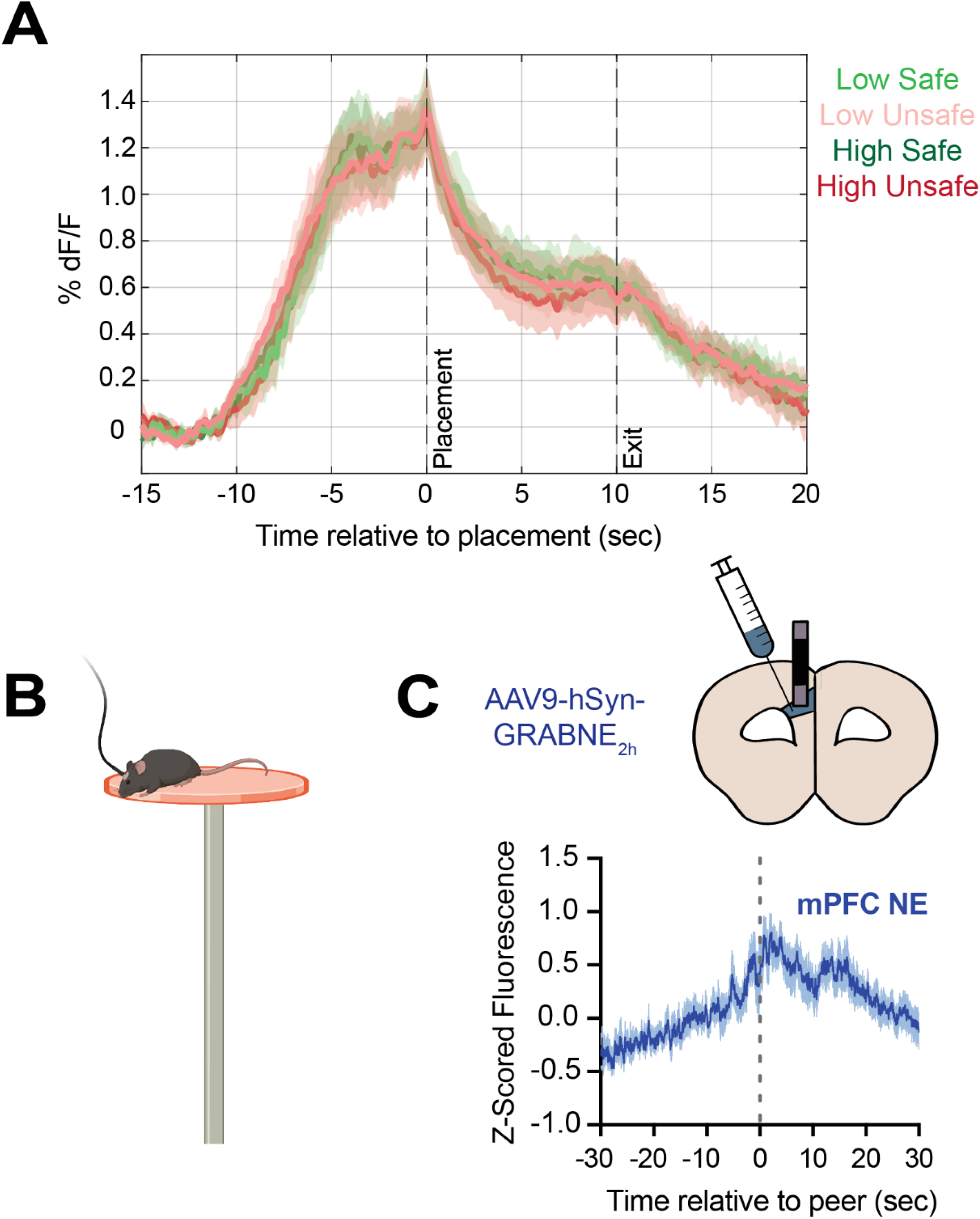
Extended characterization of prelimbic norepinephrine release to height threat. A) Plot of PL nore-pinephrine release during virtual visual cliff pole descent task across stimulus (high, low) x outcome (safe, unsafe) pairs. No differences are observed in time-warped mouse-level analyses. B) Schematic of elevated platform task. A “peer” occurs when the mouse’s head is over the edge of the platform and pointed towards the ground. C) Mouse-averaged PL norepinephrine release (measured using AAV9-hSyn-GRABNE_2h)_ aligned to the time of the peer demonstrates increased PL norepinephrine release upon the sight of heights.

**Supplemental Figure 3.**
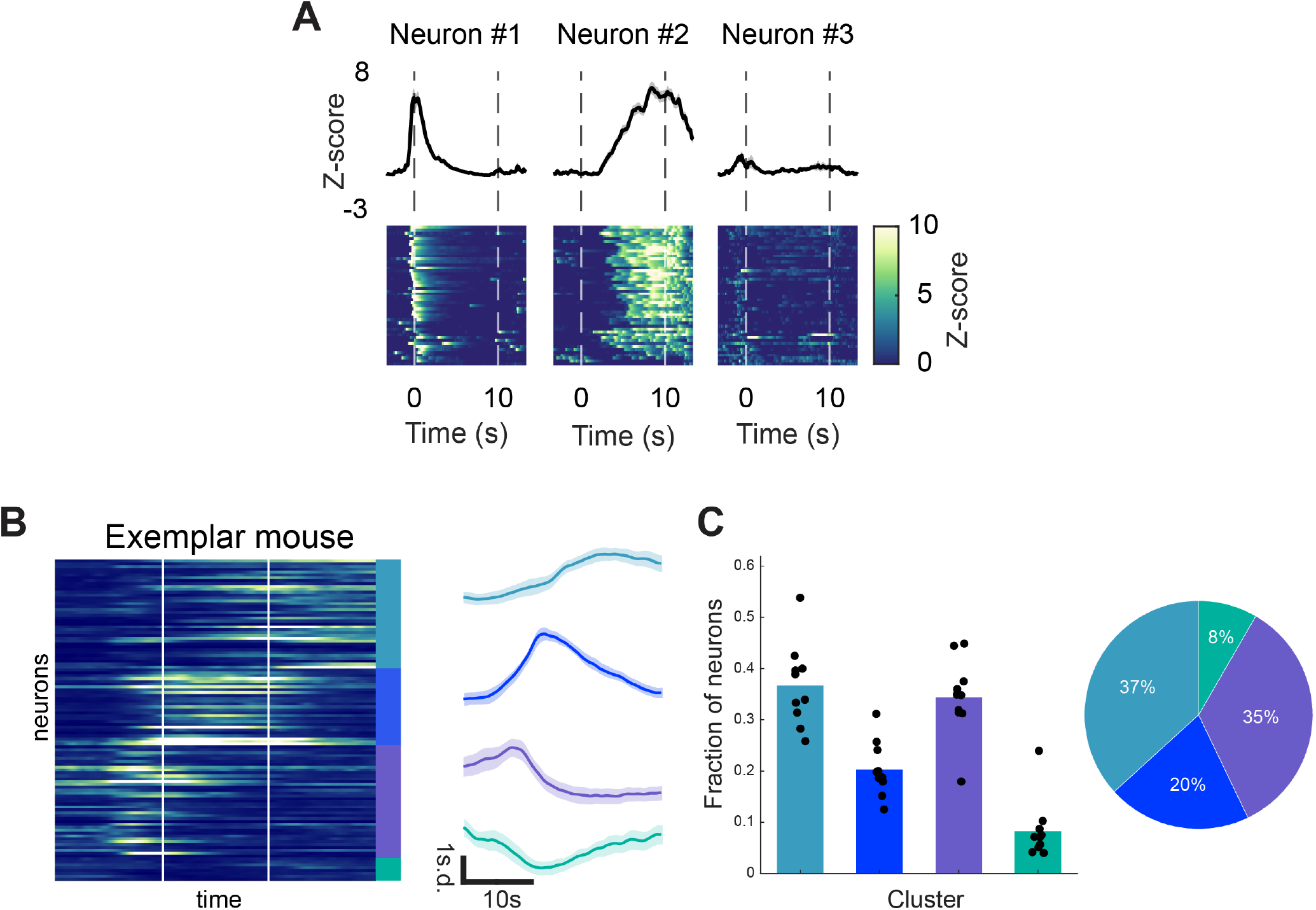
Time-warped analysis of prelimbic neural responses. A) Exemplar neurons time warped from placement to exit (Placement 0s, exit 10s). Each neuron’s average z-scored trace (top) and activity over trials (heatmaps, bottom) show similar profiles to position-based analysis. B-C) Hierarchical clustering revealed 4 temporal response profiles. B, exemplar mouse neural activity averaged over trials for neurons within each cluster. C, quantification of proportion of neurons within each cluster per mouse.

## References

1. Krystal, J. H. et al. It Is Time to Address the Crisis in the Pharmacotherapy of Posttraumatic Stress Disorder: A Consensus Statement of the PTSD Psychopharmacology Working Group. Biol Psychiatry 82, e51–e59 (2017).

2. Dunsmoor, J. E., Cisler, J. M., Fonzo, G. A., Creech, S. K. & Nemeroff, C. B. Laboratory models of post-traumatic stress disorder: The elusive bridge to translation. Neuron 110, 1754–1776 (2022).

3. Deslauriers, J., Toth, M., Der-Avakian, A. & Risbrough, V. B. Current Status of Animal Models of Posttraumatic Stress Disorder: Behavioral and Biological Phenotypes, and Future Challenges in Improving Translation. Biological Psychiatry 83, 895–907 (2018).

4. Raut, R. V. et al. Arousal as a universal embedding for spatiotemporal brain dynamics. Nature 647, 454–461 (2025).

5. Urai, A. E., Doiron, B., Leifer, A. M. & Churchland, A. K. Large-scale neural recordings call for new insights to link brain and behavior. Nat Neurosci 25, 11–19 (2022).

6. Breton-Provencher, V., Drummond, G. T. & Sur, M. Locus Coeruleus Norepinephrine in Learned Behavior: Anatomical Modularity and Spatiotemporal Integration in Targets. Front Neural Circuits 15, 638007 (2021).

7. Schwarz, L. A. & Luo, L. Organization of the locus coeruleusnorepinephrine system. Curr Biol 25, R1051–R1056 (2015).

8. Loughlin, S. E., Foote, S. L. & Bloom, F. E. Efferent projections of nucleus locus coeruleus: topographic organization of cells of origin demonstrated by three-dimensional reconstruction. Neuroscience 18, 291–306 (1986).

9. Jones, B. E. & Moore, R. Y. Ascending projections of the locus coeruleus in the rat. II. Autoradiographic study. Brain Res 127, 25–53 (1977).

10. Loughlin, S. E., Foote, S. L. & Grzanna, R. Efferent projections of nucleus locus coeruleus: morphologic subpopulations have different efferent targets. Neuroscience 18, 307–19 (1986).

11. Loughlin, S. E., Foote, S. L. & Fallon, J. H. Locus coeruleus projections to cortex: topography, morphology and collateralization. Brain Res Bull 9, 287–294 (1982).

12. Berridge, C. W. Noradrenergic modulation of arousal. Brain Res Rev 58, 1–17 (2008).

13. Breton-Provencher, V. & Sur, M. Active control of arousal by a locus coeruleus GABAergic circuit. Nat Neurosci 22, 218–228 (2019).

14. Coull, J. T. Neural correlates of attention and arousal: insights from electrophysiology, functional neuroimaging and psychopharmacology. Prog Neurobiol 55, 343–61 (1998).

15. Foote, S. L., Berridge, C. W., Adams, L. M. & Pineda, J. A. Electrophysiological evidence for the involvement of the locus coeruleus in alerting, orienting, and attending. Prog Brain Res 88, 521–532 (1991).

16. Robbins, T. W. Cortical noradrenaline, attention and arousal. Psychol Med 14, 13–21 (1984).

17. Jones, B. E. The role of noradrenergic locus coeruleus neurons and neighboring cholinergic neurons of the pontomesencephalic tegmentum in sleep-wake states. Prog Brain Res 88, 533–543 (1991).

18. Sara, S. J. & Bouret, S. Orienting and reorienting: the locus coeruleus mediates cognition through arousal. Neuron 76, 130–41 (2012).

19. Foote, S. L., Aston-Jones, G. & Bloom, F. E. Impulse activity of locus coeruleus neurons in awake rats and monkeys is a function of sensory stimulation and arousal. Proc Natl Acad Sci U S A 77, 3033–7 (1980).

20. Jacob, S. N. & Nienborg, H. Monoaminergic Neuromodulation of Sensory Processing. Front Neural Circuits 12, 51 (2018).

21. Waterhouse, B. D. & Navarra, R. L. The locus coeruleus-norepinephrine system and sensory signal processing: A historical review and current perspectives. Brain Res 1709, 1–15 (2019).

22. Waterhouse, B. D., Moises, H. C. & Woodward, D. J. Phasic activation of the locus coeruleus enhances responses of primary sensory cortical neurons to peripheral receptive field stimulation. Brain Res 790, 33–44 (1998).

23. Devilbiss, D. M., Page, M. E. & Waterhouse, B. D. Locus ceruleus regulates sensory encoding by neurons and networks in waking animals. J Neurosci 26, 9860–9872 (2006).

24. Aston-Jones, G. & Cohen, J. D. An integrative theory of locus coeruleus-norepinephrine function: adaptive gain and optimal performance. Annu Rev Neurosci 28, 403–450 (2005).

25. Díaz-Mataix, L. et al. Characterization of the amplificatory effect of norepinephrine in the acquisition of Pavlovian threat associations. Learn Mem 24, 432–439 (2017).

26. Gresack, J. E. & Risbrough, V. B. Corticotropin-releasing factor and noradrenergic signalling exert reciprocal control over startle reactivity. Int J Neuropsychopharmacol 14, 1179–1194 (2011).

27. Mason, S. T. & Fibiger, H. Noradrenaline, fear and extinction. Brain Res 165, 47–56 (1979).

28. Davis, M., Redmond, D. E. & Baraban, J. M. Noradrenergic agonists and antagonists: effects on conditioned fear as measured by the potentiated startle paradigm. Psychopharmacology (Berl) 65, 111–118 (1979).

29. Rasmussen, K. & Jacobs, B. L. Single unit activity of locus coeruleus neurons in the freely moving cat. II. Conditioning and pharmacologic studies. Brain Res 371, 335–344 (1986).

30. Glaeser-Khan, S. et al. Spatiotemporal Organization of Prefrontal Norepinephrine Influences Neuronal Activity. eNeuro 11, ENEURO.0252-23.2024 (2024).

31. Chandler, D. J.Gao, W.-J. & Waterhouse, B. D. Heterogeneous organization of the locus coeruleus projections to prefrontal and motor cortices. Proc. Natl. Acad. Sci. U.S.A. 111, 6816–6821 (2014).

32. Totah, N. K., Neves, R. M., Panzeri, S., Logothetis, N. K. & Eschenko, O. The Locus Coeruleus Is a Complex and Differentiated Neuromodulatory System. Neuron 99, 1055–1068.e6 (2018).

33. Chandler, D. J. et al. Redefining Noradrenergic Neuromodulation of Behavior: Impacts of a Modular Locus Coeruleus Architecture. J Neurosci 39, 8239–8249 (2019).

34. Uematsu, A. et al. Modular organization of the brainstem noradrenaline system coordinates opposing learning states. Nat Neurosci 20, 1602–1611 (2017).

35. Bouras, N. N., Mack, N. R. & Gao, W.-J. Prefrontal modulation of anxiety through a lens of noradrenergic signaling. Front. Syst. Neurosci. 17, 1173326 (2023).

36. Agster, K. L., Mejias-Aponte, C. A., Clark, B. D. & Waterhouse, B. D. Evidence for a regional specificity in the density and distribution of noradrenergic varicosities in rat cortex. J Comp Neurol 521, 2195–207 (2013).

37. Cerpa, J.-C., Marchand, A. R. & Coutureau, E. Distinct regional patterns in noradrenergic innervation of the rat prefrontal cortex. Journal of Chemical Neuroanatomy 96, 102–109 (2019).

38. Burgos-Robles, A., Vidal-Gonzalez, I. & Quirk, G. J. Sustained conditioned responses in prelimbic prefrontal neurons are correlated with fear expression and extinction failure. J Neurosci 29, 8474–8482 (2009).

39. Corcoran, K. A. & Quirk, G. J. Activity in prelimbic cortex is necessary for the expression of learned, but not innate, fears. J Neurosci 27, 840–844 (2007).

40. Vidal-Gonzalez, I., Vidal-Gonzalez, B., Rauch, S. L. & Quirk, G. J. Microstimulation reveals opposing influences of prelimbic and infralimbic cortex on the expression of conditioned fear. Learn Mem 13, 728–733 (2006).

41. Burgos-Robles, A. et al. Amygdala inputs to prefrontal cortex guide behavior amid conflicting cues of reward and punishment. Nat Neurosci 20, 824–835 (2017).

42. Friedman, A. et al. A Corticostriatal Path Targeting Striosomes Controls Decision-Making under Conflict. Cell 161, 1320–1333 (2015).

43. Itzhak, Y., Perez-Lanza, D. & Liddie, S. The strength of aversive and appetitive associations and maladaptive behaviors. IUBMB Life 66, 559–571 (2014).

44. Yilmaz, M. & Meister, M. Rapid innate defensive responses of mice to looming visual stimuli. Curr Biol 23, 2011–5 (2013).

45. Cain, C. K. Beyond Fear, Extinction, and Freezing: Strategies for Improving the Translational Value of Animal Conditioning Research. Curr Top Behav Neurosci 64, 19–57 (2023).

46. Blanchard, D. C., Blanchard, R. J. & Griebel, G. Defensive responses to predator threat in the rat and mouse. Curr Protoc Neurosci Chapter 8, Unit 8.19 (2005).

47. Bastos, A. F. et al. Stop or move: Defensive strategies in humans. Behavioural Brain Research 302, 252–262 (2016).

48. Mobbs, D. et al. When Fear Is Near: Threat Imminence Elicits Prefrontal-Periaqueductal Gray Shifts in Humans. Science 317, 1079–1083 (2007).

49. Caroline Blanchard, D., Hynd, A. L., Minke, K. A., Minemoto, T. & Blanchard, R. J. Human defensive behaviors to threat scenarios show parallels to fear- and anxiety-related defense patterns of non-human mammals. Neuroscience & Biobehavioral Reviews 25, 761–770 (2001).

50. Bertram, T. et al. Human threat circuits: Threats of pain, aggressive conspecific, and predator elicit distinct BOLD activations in the amygdala and hypothalamus. Front. Psychiatry 13, 1063238 (2023).

51. Ren, C. & Tao, Q. Neural Circuits Underlying Innate Fear. Adv Exp Med Biol 1284, 1–7 (2020).

52. Gibson, E. J. & Walk, R. D. The ‘visual cliff’. Sci Am 202, 64–71 (1960).

53. Bauer, J. H. Development of visual cliff discrimination by infant hooded rats. J Comp Physiol Psychol 84, 380–5 (1973).

54. Fox, M. W. The visual cliff test for the study of visual depth perception in the mouse. Anim Behav 13, 232–3 (1965).

55. Schwartz, A. N., Campos, J. J. & Baisel, E. J. The visual cliff: cardiac and behavioral responses on the deep and shallow sides at five and nine months of age. J Exp Child Psychol 15, 86–99 (1973).

56. Yilmaz Balban, M. et al. Human Responses to Visually Evoked Threat. Curr Biol 31, 601–612 e3 (2021).

57. Boone, H. C. et al. Natural binocular depth discrimination behavior in mice explained by visual cortical activity. Curr Biol 31, 2191–2198 e3 (2021).

58. Liu, J., Lin, L. & Wang, D. V. Representation of Fear of Heights by Basolateral Amygdala Neurons. J Neurosci 41, 1080–1091 (2021).

59. Shang, W. et al. A non-image-forming visual circuit mediates the innate fear of heights in male mice. Nat Commun 15, 3746 (2024).

60. Zhu, H. Y., Chen, H. T. & Lin, C. T. The Effects of Virtual and Physical Elevation on Physiological Stress during Virtual Reality Height Exposure. IEEE Trans Vis Comput Graph PP, (2021).

61. Newman, V. E., Liddell, B. J., Beesley, T. & Most, S. B. Failures of executive function when at a height: Negative height-related appraisals are associated with poor executive function during a virtual height stressor. Acta Psychol (Amst) 203, 102984 (2020).

62. Raffegeau, T. E. et al. Walking (and talking) the plank: dual-task performance costs in a virtual balance-threatening environment. Exp Brain Res 242, 1237–1250 (2024).

63. Drummond, G. T. et al. Cortical astrocytes extend phasic norepinephrine signals and critically mediate learned behavior. Preprint at 10.1101/2024.10.24.620009 (2024).

64. Liu, X. et al. Neural circuit underlying individual differences in visual escape habituation. Neuron 113, 2344–2357.e5 (2025).

65. Trent, S. et al. Fear conditioning: Insights into learning, memory and extinction and its relevance to clinical disorders. Progress in NeuroPsychopharmacology and Biological Psychiatry 138, 111310 (2025).

66. Quirk, G. J., Repa, J. C. & LeDoux, J. E. Fear conditioning enhances short-latency auditory responses of lateral amygdala neurons: Parallel recordings in the freely behaving rat. Neuron 15, 1029–1039 (1995).

67. Resulaj, A., Kiani, R., Wolpert, D. M. & Shadlen, M. N. Changes of mind in decision-making. Nature 461, 263–266 (2009).

68. Lak, A. et al. Reinforcement biases subsequent perceptual decisions when confidence is low, a widespread behavioral phenomenon. eLife 9, e49834 (2020).

69. Ashwood, Z. C. et al. Mice alternate between discrete strategies during perceptual decision-making. Nat Neurosci 25, 201–212 (2022).

70. Rabbitt, P. M. Errors and error correction in choice-response tasks. Journal of Experimental Psychology 71, 264–272 (1966).

71. Narayanan, N. S., Cavanagh, J. F., Frank, M. J. & Laubach, M. Common medial frontal mechanisms of adaptive control in humans and rodents. Nat Neurosci 16, 1888–1895 (2013).

72. Nieuwenhuis, S., Aston-Jones, G. & Cohen, J. D. Decision making, the P3, and the locus coeruleus–norepinephrine system. Psychological Bulletin 131, 510–532 (2005).

73. Yerkes, R. M. & Dodson, J. D. The relation of strength of stimulus to rapidity of habit formation. Journal of Comparative Neurology and Psychology 18, 459–482 (1908).

74. Mahn, M. et al. High-efficiency optogenetic silencing with soma-targeted anion-conducting channelrhodopsins. Nat Commun 9, 4125 (2018).

75. Williams, A. H. et al. Discovering Precise Temporal Patterns in Large-Scale Neural Recordings through Robust and Interpretable Time Warping. Neuron 105, 246–259.e8 (2020).

76. Basu, A. et al. Frontal norepinephrine represents a threat prediction error under uncertainty. Biol Psychiatry 96, 256–267 (2024).

77. Feng, J. et al. A Genetically Encoded Fluorescent Sensor for Rapid and Specific In Vivo Detection of Norepinephrine. Neuron 102, 745–761 e8 (2019).

78. Feng, J. et al. Monitoring norepinephrine release in vivo using next-generation GRABNE sensors. Neuron 112, 1930–1942.e6 (2024).

79. Carion, N. et al. SAM 3: Segment Anything with Concepts. Preprint at 10.48550/ARXIV.2511.16719 (2025).

80. Barbosa, J. et al. Quantifying Differences in Neural Population Activity With Shape Metrics. Preprint at 10.1101/2025.01.10.632411 (2025).

81. Battivelli, D., Fan, Z., Hu, H. & Gross, C. T. How can ethology inform the neuroscience of fear, aggression and dominance? Nat. Rev. Neurosci. 25, 809–819 (2024).

82. Dennis, E. J. et al. Systems Neuroscience of Natural Behaviors in Rodents. J Neurosci 41, 911–919 (2021).

83. Miller, C. T. et al. Natural behavior is the language of the brain. Curr Biol 32, R482–R493 (2022).

84. Michaiel, A. M., Abe, E. T. & Niell, C. M. Dynamics of gaze control during prey capture in freely moving mice. Elife 9, (2020).

85. Parker, P. R. L. et al. Distance estimation from monocular cues in an ethological visuomotor task. Elife 11, (2022).

86. Krakauer, J. W., Ghazanfar, A. A., Gomez-Marin, A., MacIver, M. A. & Poeppel, D. Neuroscience Needs Behavior: Correcting a Reductionist Bias. Neuron 93, 480–490 (2017).

87. Kirschner, H., Humann, J., Derrfuss, J., Danielmeier, C. & Ullsperger, M. Neural and behavioral traces of error awareness. Cogn Affect Behav Neurosci 21, 573–591 (2021).

88. Ako, R.Terada, S.-I. & Matsuzaki, M. A cerebello-thalamo-cortical pathway transmits reward-based post-error signals for motor timing correction during learning in male mice. Nat Commun 16, 7663 (2025).

89. Norman, K. J. et al. Post-error recruitment of frontal sensory cortical projections promotes attention in mice. Neuron 109, 1202–1213.e5 (2021).

90. Mansouri, F. A. et al. Direct current stimulation of prefrontal cortex modulates error-induced behavioral adjustments. Eur J of Neuroscience 44, 1856–1869 (2016).

91. Fu, Z. et al. Single-Neuron Correlates of Error Monitoring and Post-Error Adjustments in Human Medial Frontal Cortex. Neuron 101, 165–177.e5 (2019).

92. Javanbakht, A. et al. The First Use of Advanced Augmented Reality Technology for Treatment of PTSD. Annals of Clinical Psychiatry 36, 155–156 (2025).

93. Breton-Provencher, V., Drummond, G. T., Feng, J., Li, Y. & Sur, M. Spa-tiotemporal dynamics of noradrenaline during learned behaviour. Na-ture 606, 732–738 (2022).

94. Rigotti, M. et al. The importance of mixed selectivity in complex cog-nitive tasks. Nature 497, 585–590 (2013).

95. Wójcik, M. J. et al. Learning shapes neural geometry in the primate prefrontal cortex. Nat Neurosci 29, 1966–1975 (2026).

96. Guidera, J. A. et al. Regional specialization in prefrontal cortex manifests in the reliability of task progression codes. Neuron 114, 1152–1161.e8 (2026).

97. Matarasso, A. et al. Stimulus-Dependent Dopamine Dynamics from Locus Coeruleus Axons. Preprint at 10.1101/2025.09.15.676390 (2025).

98. Kempadoo, K. A., Mosharov, E. V., Choi, S. J., Sulzer, D. & Kandel, E. R. Dopamine release from the locus coeruleus to the dorsal hippocampus promotes spatial learning and memory. Proc. Natl. Acad. Sci. U.S.A. 113, 14835–14840 (2016).

99. Devoto, P., Flore, G., Saba, P., Fà, M. & Gessa, G. L. Stimulation of the locus coeruleus elicits noradrenaline and dopamine release in the medial prefrontal and parietal cortex. Journal of Neurochemistry 92, 368–374 (2005).

100. Yang, J.-H. et al. Frontal cortex norepinephrine, serotonin, and dopamine dynamics in an innate fear-reward behavioral model. Preprint at 10.1101/2023.11.27.568929 (2023).

101. Simpson, E. H. et al. Lights, fiber, action! A primer on in vivo fiber photometry. Neuron 112, 718–739 (2024).

102. McCall, J. G. et al. Locus coeruleus to basolateral amygdala noradrenergic projections promote anxiety-like behavior. eLife 6, e18247 (2017).

103. Pnevmatikakis, E. A. & Giovannucci, A. NoRMCorre: An online algorithm for piecewise rigid motion correction of calcium imaging data. J Neurosci Methods 291, 83–94 (2017).

104. Pnevmatikakis, E. A. et al. Simultaneous Denoising, Deconvolution, and Demixing of Calcium Imaging Data. Neuron 89, 285–99 (2016).

